# 3D Hydrodynamic Flow Lithography

**DOI:** 10.1101/2025.10.27.684743

**Authors:** Yiying Zou, Mehmet Akif Sahin, Muhammad Zia Ullah Khan, Muhammad Usman Akhtar, Daniel Stoecklein, Ghulam Destgeer

## Abstract

Continuous- and stop-flow lithography has been widely used for the fabrication of ingenious multi-dimensional microstructures, including fibers and microparticles. The flow profile of multiple co-flowing streams during the process, as one of the dimensions, dictates the cross-sectional morphology of the microstructures. Here, we introduce a three-dimensional hydrodynamic flow lithography (3D HFL) approach that enables rapid and programmable generation of desired flow profiles and their conversion into solid microstructures. By strategically combining multiple flow sculpting strategies, including variable inlet configurations, intra-channel pillar configurations, and outlet configurations, we achieved precise and versatile control over flow sculpting, as validated by computational fluid dynamics simulations and experimental production of microparticles and fibers, offering unparalleled design flexibility. Importantly, the flow sculpting device is fabricated using low-cost 3D-printed molds to cast PDMS channels with precisely aligned inlets and complex geometries that are difficult or impossible to achieve using conventional soft lithography. Our 3D HFL approach effectively overcomes the limitations of existing microfabrication techniques in device complexity, structural diversity, multi-material integration, and production throughput. Furthermore, we demonstrated the capability of this platform to produce anisotropic multi-material microparticles and fibers tailored for specific applications, such as amphiphilic particles for uniform microdroplet capture, biocompatible patterned hydrogels for controlled cell adhesion, and dual-layer fibers for temperature sensing. The simplicity of device fabrication, combined with the broad design flexibility, establishes this platform as a scalable, high-throughput, and versatile solution for engineering anisotropic microparticles and fibers with tailored functionalities, providing powerful tools for a wide range of downstream fields.

## 1. Introduction

The functionalities of microstructures, e.g., microparticles and fibers, are fundamentally governed by their size, structure, and composition[1–4]. These physical attributes enable a wide range of specialized uses. For example, shape- or color-barcoded microparticles have been successfully employed as multiplexed bioassay platforms for allowing simultaneous detection of multiple analytes[5–7]. In parallel, biocompatible spherical or crescent-shaped particles[8–10], as well as microfibers[11], have emerged as ideal scaffolds for culture and analysis, providing well-controlled microenvironments for cell-level studies. Beyond static applications, microparticles also serve dynamic roles in micro-robotics. The magnetic, optical, and acoustic actuation capabilities of micro-robots typically stem from either responsive material compositions[12, 13] or tailored geometries[14, 15], which facilitate deformation, rotation, or translational motion.

These diverse applications underscore the critical need for fabrication techniques capable of reliably producing functional microparticles and fibers with precisely tunable sizes, shapes, and compositions. Comprising continuous-flow lithography (CFL)[16] and stop-flow lithography (SFL)[17], flow lithography (FL)[18] has emerged as a particularly versatile technique for generating these microstructures with a wide range of functionalities. Early implementations of SFL allowed the creation of 2D microparticles through the modulation of 2D photomask patterns, yielding geometrically simple but functionally robust architecture[19, 20]. Innovations such as phase-mask interference[21], two-step exposure[22], non-uniform UV exposure or absorption[23–25], and intra-channel flow restrictions[26] expanded this capability to 2.5D structures with controlled depth-wise heterogeneity. Additional approaches, including soft membrane[27] and vertical[28] flow lithography, have pushed the boundaries toward true 3D structuring. However, these methods often depend on precise, multi-step photomask alignment and are typically limited by moderate throughput and compositional flexibility.

An alternative for fabricating 3D structured microparticles emerges in intersecting the 2D UV pattern projected from the photomask with an engineered multi-layered flow profile within microfluidic channels. Pioneering work by the Doyle group established hydrodynamic focusing lithography[29], which employed bilayer polydimethylsiloxane (PDMS) microchannels to generate a dual-axis, multi-layered flow profile. However, despite its strengths, the technique is constrained by its reliance on relatively basic channel geometries and intricate alignment steps for multilayered microchannel fabrication, making it challenging to produce more sophisticated flow profile designs. Subsequently, the Di Carlo group developed an inertial flow sculpting method[30], which employs single-layered PDMS microchannels embedded with pillar structures to generate secondary Dean flows at high Reynolds (*Re*) numbers, thereby dramatically expanding the diversity of achievable flow profile[31–36]. However, the need for high-pressure inertial flow at *Re ∼O*(10) or higher, along with extended pillar sequences within long microfluidic channels to achieve the desired flow profiles, can pose significant challenges, such as increased channel capacitance and long flow stopping times under high inlet pressure, and lower material yield due to higher precursor fluid waste between SFL cycles caused by high flow rates. Nonetheless, both inertial flow sculpting[30–36] and coaxial 3D-printed lithography represent meaningful advances over the hydrodynamic focusing approach[29]. More recently, we have demonstrated 3D-printed microfluidic devices with customizable nozzles to sculpt a coaxial flow profile at low *Re ∼O*(1) or smaller to produce a wide variety of shape-encoded, multi-material particles[6, 37–40]. This approach expands the design space through the flexibility of 3D printing but relies on oxygen-impermeable channels, necessitating inert sheath flows to prevent clogging, which is not an issue with conventional PDMS-based microchannels used in hydrodynamic and inertial flow sculpting methods. We summarize the features of conventional SFL-based particle fabrication techniques for comparison in the supporting information (Table S1). To address these trade-offs, we envision a hybrid strategy that combines the oxygen permeability of PDMS microchannels with the design flexibility of 3D printing, aiming to enable scalable, sheath-free fabrication of complex microparticles and fibers[41] with enhanced structural tunability and functional versatility. The 3D HFL platform introduced here addresses specific limitations that none of the prior methods resolve simultaneously: unlike hydrodynamic focusing lithography[29], it does not require bilayer PDMS fabrication or manual port alignment; unlike inertial flow sculpting[30–36], it operates at *Re* ≲ 1 and therefore avoids the long flow-stopping times and high precursor waste associated with high-pressure operation; and unlike coaxial 3D-printed lithography[6, 37–40], it uses oxygen-permeable PDMS channels and requires no inert sheath flow, directly enabling closed cross-sectional geometries (e.g., O-shaped profiles). Together, these distinctions—comprehensively tabulated in Table S1—motivate the hybrid 3D-printed mould plus soft-lithography strategy adopted here.

In this work, we present a 3D hydrodynamic flow lithography (3D HFL) method (Figure 1) for producing multi-material, anisotropic microparticles and fibers with engineered shapes and tunable sizes. We use an oxygen-permeable 3D PDMS microfluidic channel with multiple inlets (up to five) converging into a single outlet (∼100–500 μm) to photo-cure particles under a cyclic UV exposure through a photomask (SFL, >10^4^ particles/hour) or long fibers under a continuous bulk UV exposure (CFL), without the need for an inert sheath flow stream (Figure 1A). For a precise positioning of the inlet ports within the main channel, we devised a new cleanroom-free microchannel fabrication workflow. We 3D-printed the low-cost channel molds with integrated pillars that form the inlet ports within the PDMS channel after the mold replica step. This hybrid device fabrication method, combining 3D printing and soft lithography, enables us to produce microfluidic devices with various inlets, pillars, and outlet configurations to shape multi-layered co-flowing streams into a wide variety of distinct flow profiles, with potential for further expansion. To predict and optimize these sculpted flow profiles, we utilized computational fluid dynamics (CFD) simulations to evaluate each channel design prior to device fabrication (Figure 1B). Subsequent experiments with the fabricated devices yielded structured particles and fibers showing multiple precursor streams cured into the desired cross-sectional shapes and compositions, demonstrating excellent agreement with our CFD predictions (Figure 1C).

**Figure 1.**
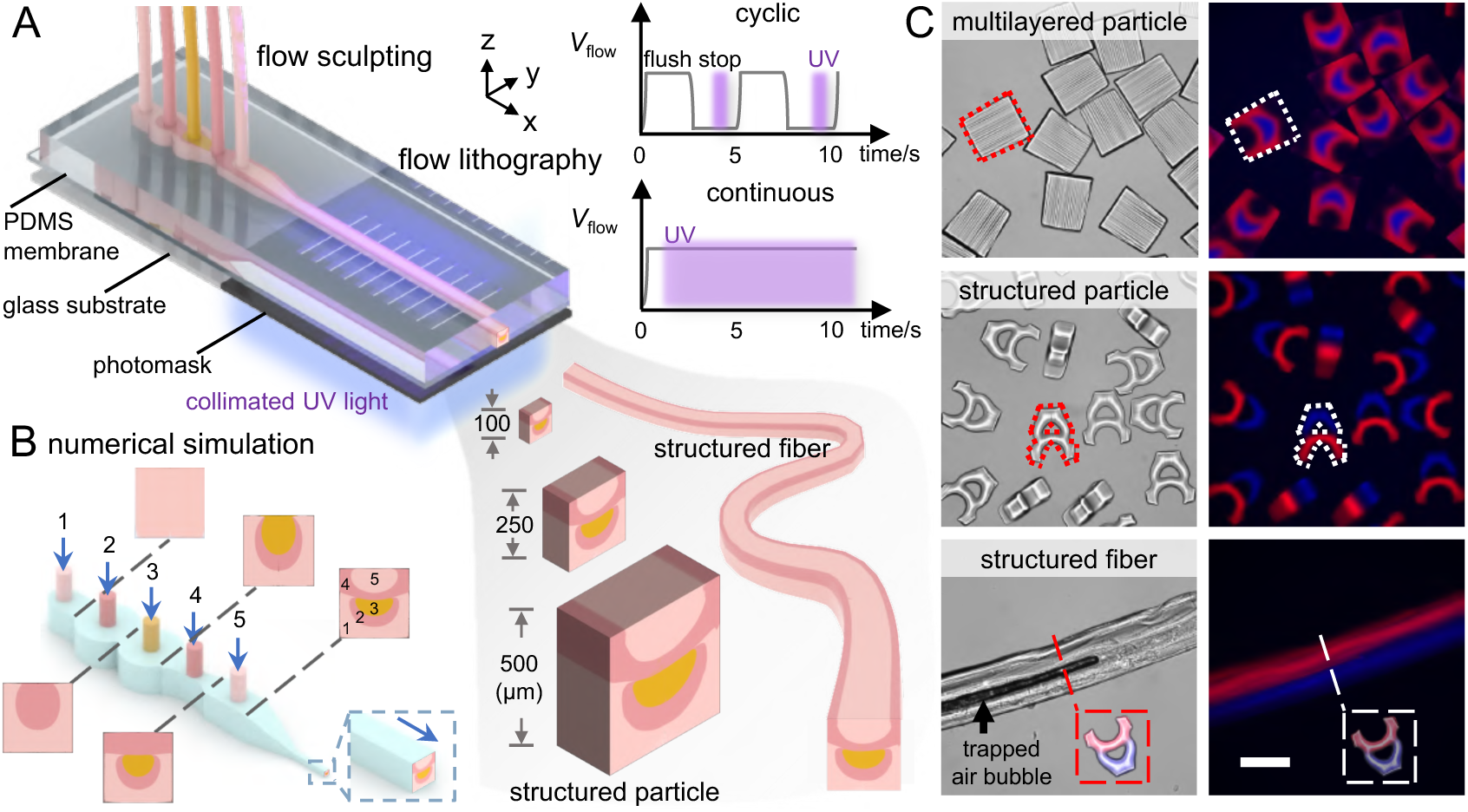
3D hydrodynamic flow lithography for producing structured microparticles and fibers. (A) A 3D hydrodynamic flow sculpting device producing multi-layered structured microparticles and fibers upon cyclic and continuous UV exposure of curable polymer precursors, respectively. Square PDMS microchannels with nominal outer dimensions of 100, 250, and 500 µm defined the outer dimensional limit of the cured particles and fibers. (B) Numerical modeling of the sculpted flow profile with five layers (1-5) controlled by the flow rate through the five independent inlets. (C) Brightfield and fluorescent microscopy images of the multilayered structured microparticles and fibers. Top: Five curable streams, transparent (1), red (2), blue (3), red (4), and transparent (5), led to multilayered particles with outer trapezoidal shape matching the channel cross-section. Middle: Three inert (1, 3, 5) and two curable blue (2) and red (4) streams resulted in bi-layered structured particles shaped like the letter “A”. Bottom: A continuous UV exposure of a similar flow profile, shaped like the letter “A”, produces a structure fiber with a matching cross-section (inset). A bubble trapped within the fiber confirms a hollow cavity.

## 2. Results and discussion

### 2.1 Flow sculpting strategies

To create structured flow profiles inside the microchannels, we have demonstrated three flow sculpting strategies using variable inlet, intra-channel pillar, and outlet configurations. First, the inlets introduce multiple streams simultaneously or sequentially within the channel at desired locations to form an initial multi-layered flow profile, which then would be further reshaped by the different indented pillar structures inside the main channel. Lastly, the cross-sectional shape and size of the outlet would determine the final cross-sectional outline of the sculpted flow before UV curing. Contrary to the conventional methods that require complex multi-layered PDMS microchannels with manually punched inlet ports and limited flow sculpting possibilities[29, 42], we leverage 3D-printed molds methodology to enable highly flexible design and easy fabrication of PDMS microchannels with precisely aligned inlet ports and pillars within the main channel as well as with variable outlet channel cross-sections, allowing multiple streams to be freely stacked and sculpted under these three flow sculpting strategies. Notably, both inertial and non-inertial regimes can theoretically be used for flow sculpting, and the same channel geometry may generate distinct flow profiles depending on the flow regime (Figure S1). In this work, we focused exclusively on the non-inertial regime and investigated the flow sculpting effects generated by different channel configurations under Stokes flow condition (*Re* ≲ 1), where *Re* is the Reynolds number. Throughout, we first characterize each sculpting strategy through numerical investigation using CFD simulations (COMSOL Multiphysics; see section 3.1), and then validate key predictions experimentally. To visualize these sculpted flow profiles, we deliberately use thin, geometrically simple square cross-sections as a controlled test bed in which the sculpted profile can be read out unambiguously and quantified using image-based metrics; the cross-sectional size and thickness control of these reference structures is characterized in Figure S6 and Note S6, and the same inlet/pillar/outlet operators introduced in Figures 2-4 are then used directly in the multi-layered architectures shown in Figure 5.

**Figure 2.**
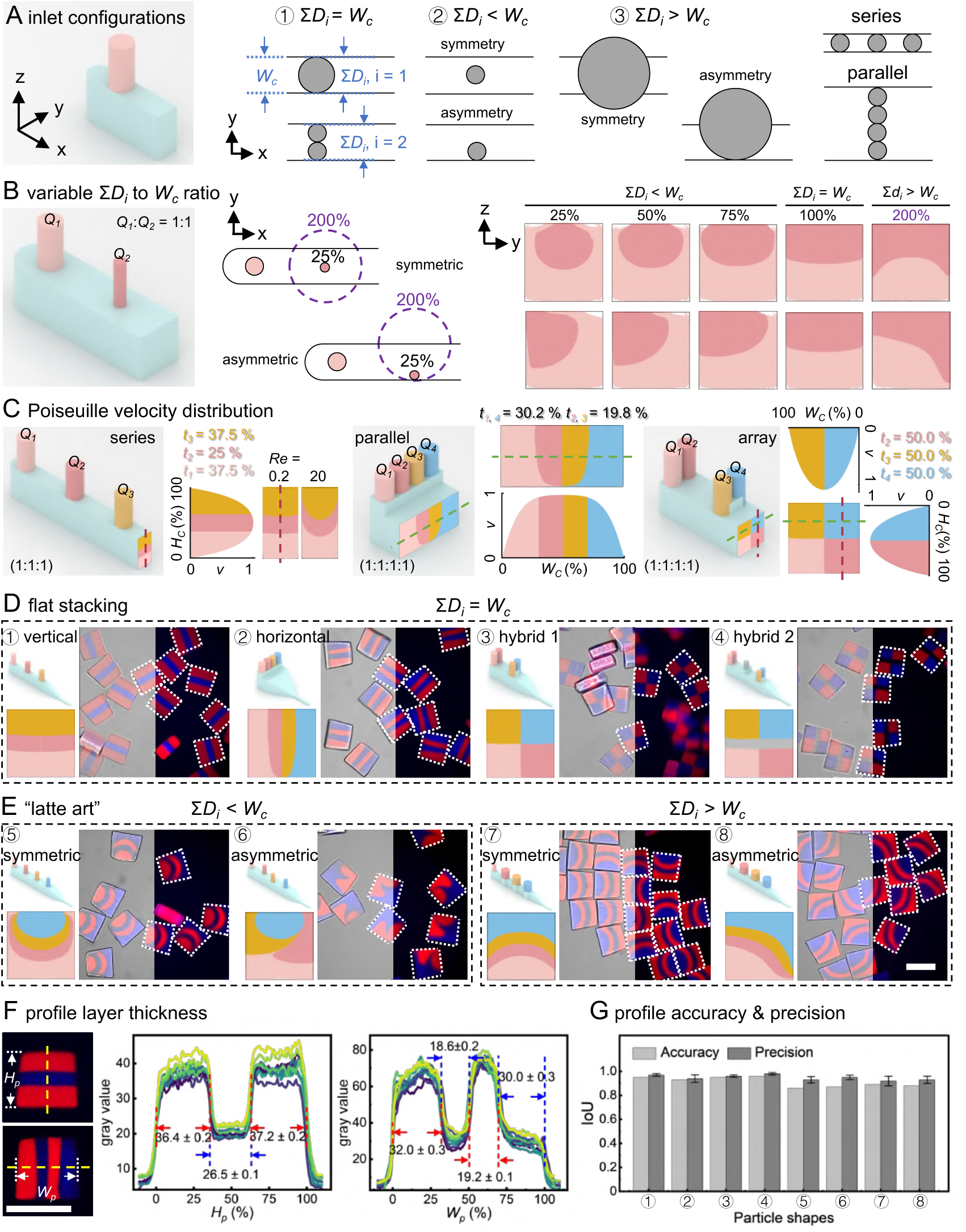
Flow sculpting using variable inlet configurations. (A) Three different inlet configurations are realized such that the sum of inlet diameters (*D_i_*) at a given YZ-plane is equal to, smaller than, or larger than the channel width (*W_c_*), i.e., Σ*D_i_* = *W_c_*, Σ*D_i_* < *W_c_*, or Σ*D_i_* > *W_c_*, respectively. The inlets can be further arranged along the X- or Y-axis in series, or in parallel configuration, respectively, within the main channel. (B) After the first inlet, any subsequent inlet can be positioned in a symmetric or asymmetric configuration with respect to the main channel. The relative diameter *D_i_* of the second inlet, i.e., 25-200% of *W_c_*, and its position with respect to the channel center lead to different sculpted cross-sectional flow profiles. Here, we show the second inlet in the center of the channel (symmetric) or tangent to a side wall (asymmetric). (C) The series and parallel inlet configurations produce vertical and horizontal stacking of the flow streams within the channel cross section. An array configuration leads to a hybrid checkered flow profile. The thicknesses of individual layers are defined by the flow rates and Poiseuille velocity profile. (D, E) The multi-layered flow streams are converged into a smaller outlet channel (250 μm × 250 μm) where structured microparticles are UV-cured. (D) The multi-layered particles are cured with vertical, horizontal, and hybrid layer stacking for Σ*D_i_* = *W_c_*. (E) Particle shapes resembling “latte art” are cured using symmetric and asymmetric configurations for Σ*D_i_* < *W_c_*, and for Σ*D_i_* > *W_c_*. (F) Dimensional characterization shows high uniformity across particles and individual layers. The layer thicknesses match the numerical predictions in (C). (G) Flow pattern accuracy (comparing simulation to experiment) and precision (similarity within a batch of microparticles, a sample size of n ≥ 10 particles was used) using intersection-over-union (IoU) scores. Scale bars: 250 μm.

**Figure 3.**
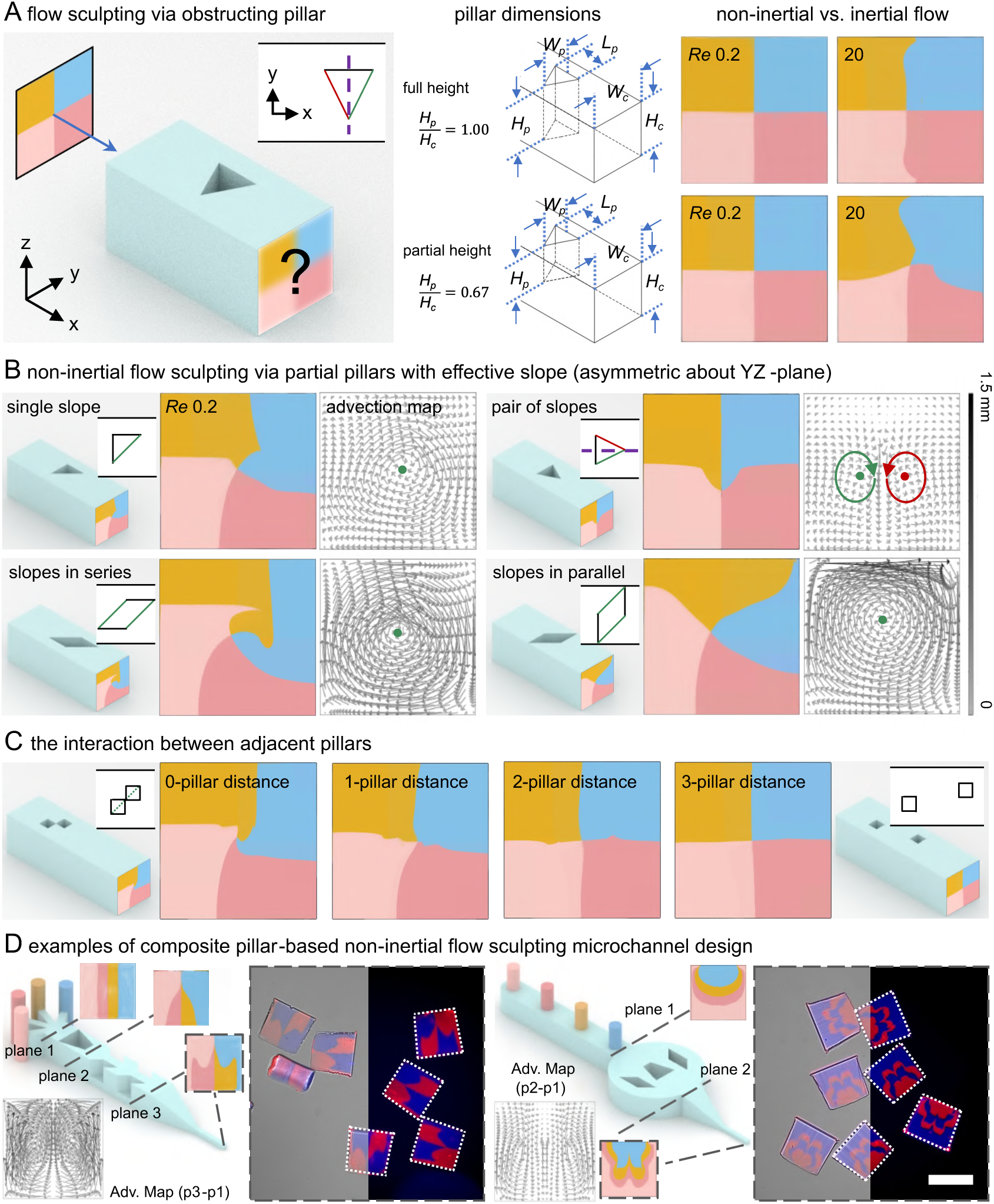
Flow sculpting using variable pillar configurations. (A) A representative flow sculpting mode demonstrates that a given initial flow profile (input) is going to be transformed into a resulting flow profile (output) after passing through an indented pillar structure (unit operator). Full-height and partial-height pillars exhibit distinct flow sculpting behaviors in non-inertial (*Re* = 0.2) and inertial (*Re* = 20) flow regimes. For pillar structures symmetric about the YZ-plane, a net-zero flow sculpting effect is produced in non-inertial flow. (B) A non-net-zero flow sculpting effect is produced in non-inertial flow only when the pillar structures are partially indented and asymmetric about the YZ-plane, regardless of whether they maintain XZ-plane symmetry or exhibit complete asymmetry. (C) Effect of the interaction between adjacent pillars on flow sculpting. (D) Representative examples of flow sculpting microchannels incorporating composite partial-pillar designs in a non-inertial flow, and fabricated uniquely shaped particles. Scale bar: 250 μm.

**Figure 4.**
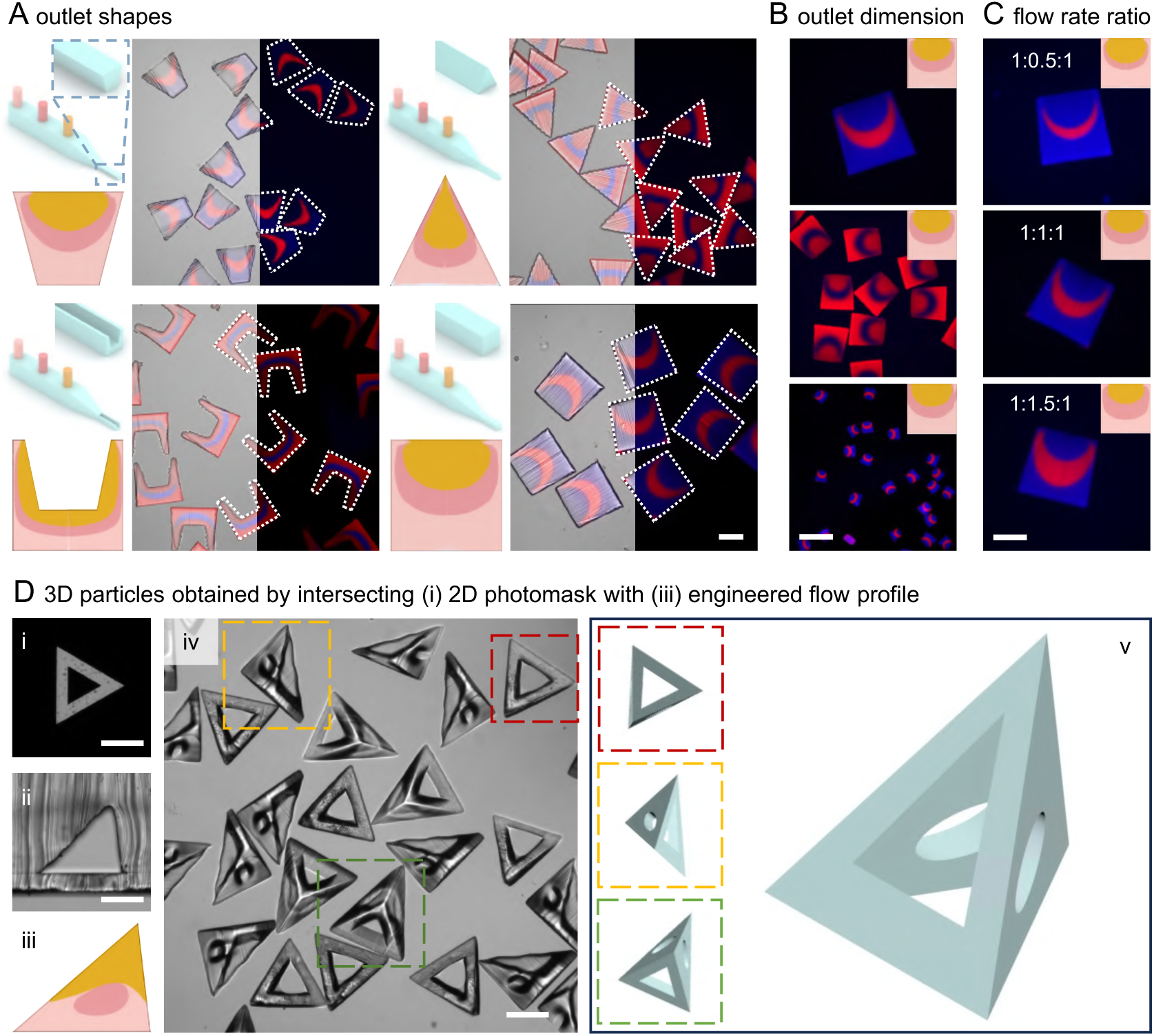
Flow sculpting using variable outlet channel configurations. Microchannel designs simulated flow profiles and experimental microparticle productions with (A) various outlet channel geometries: inverted trapezoidal, triangular, slit-shaped, and square outlines. (B) Square outlet channel geometry with various dimensions: ∼ 500 μm, 250 μm, and 100 μm side length. (C) 500 μm square-shaped multi-layered microparticles were produced under various pressure ratios between the three inlets. (D) (i) the 2D photomask projected onto (ii) the structured microfluidic channel containing (iii) an engineered multi-layered flow profile, cured into (iv) the resulting particles, shown with (v) the corresponding 3D schematic. Scale bars: 250 μm.

**Figure 5.**
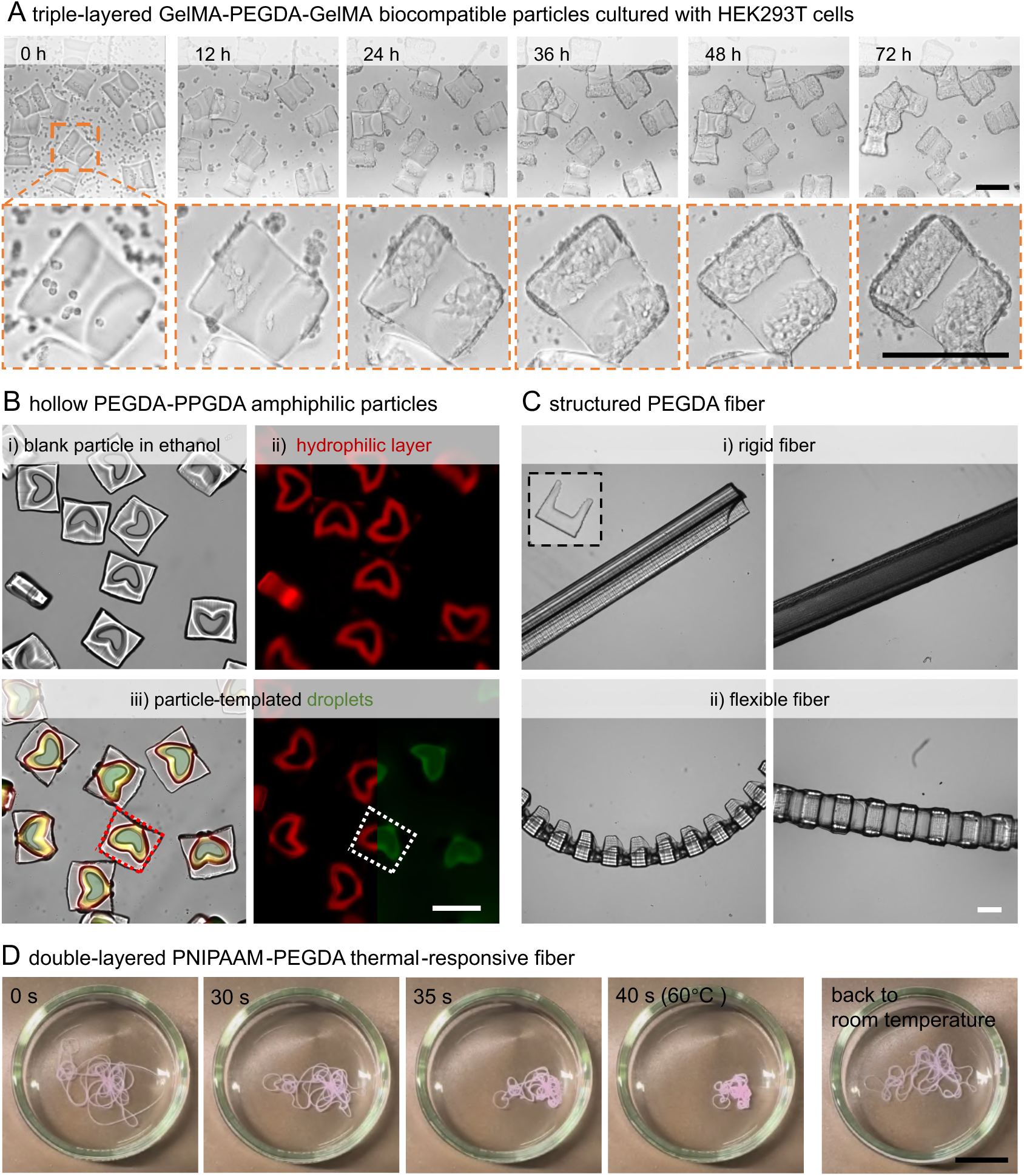
Representative microparticles and fibers fabricated via 3D hydrodynamic flow lithography. (A) Time-lapse images showing targeted cell adhesion on patterned, biocompatible PEGDA**-**GelMA microparticles. (B) Microfluidics-free droplet formation using hollow, amphiphilic PEGDA**-**PPGDA microparticles. (C) Slit-shaped PEGDA microparticles and fibers with anisotropic features along the fiber length. (D) Thermal-responsive PEGDA-PNIPAAM microfibers. Scale bars: 250 μm for (A)–(C), 1 cm for (D).

#### 2.1.1 Flow sculpting using variable inlet configurations

Using CFD simulations, we investigated three different inlet configurations to sculpt multiple co-flowing streams (Figure 2A). In the first inlet configuration, the diameter (*D_i_*) of a single circular inlet is equal to the channel width (*W_c_*), however, for multiple adjacent inlets placed at a single YZ-plane, the sum of inlet diameters is still equal to *W_c_*, i.e., Σ*D_i_* = *W_c_*. In the second inlet configuration, an inlet with a relatively smaller diameter with respect to the channel width is placed symmetrically in the middle of the channel or asymmetrically closer to one of the side walls, such that Σ*D_i_* < *W_c_*. Lastly, in the third inlet configuration, an inlet with a diameter larger than the channel width, i.e., Σ*D_i_* > *W_c_*, is placed symmetrically or asymmetrically over the microchannel width. Multiple inlets, each with any of the three inlet configurations, can be further arranged in series or in parallel formation within the main channel along the X- or Y-axis, respectively.

The configuration of the first inlet in the fluidic circuit with *i* = 1 does not have any significance, as it simply fills the channel with an initial fluid, and secondary flows are diminished within a channel width; however, the size and position of any subsequent inlet will have a significant effect on the sculpting of a multi-layered flow profile (Figure 2B). For example, the relative diameter *D_i_* of the second inlet varied as a fraction of the channel width *W_c_*, i.e., *D_i_* = 25-200% of *W_c_*, and its position with respect to the channel center, i.e., a symmetric or asymmetric configuration, can lead to different sculpted cross-sectional flow profiles. For configuration one with Σ*D_i_* = *W_c_*, a square microchannel having two inlets predicts a perfect vertical stacking of two flow streams. For Σ*D_i_* < *W_c_*, the flow stream from the second inlet forms an upward curved layer on top, where the curvature of the interface and the bi-layer cross-sectional flow profile are determined by the relative inlet diameter *D_i_* with respect to the channel width *W_c_*, i.e., 25, 50, or 75%, and the relative position of the inlet within the channel. For Σ*D_i_* > *W_c_*, the second flow stream forms an oppositely curved interface, further tuned by the symmetric or asymmetric positioning of the inlet.

In a multi-layered sculpted flow profile, the thicknesses and spatial positionings of individual layers within the channel cross-section are governed by the flow rate ratio and the selected inlet configuration, respectively (Figure 2C). For example, an inlet configuration (Σ*D_i_* = *W_c_*) with three inlets connected to the main channel in series produces a vertical stacking of three flow streams within the channel cross-section such that a flow rate ratio *Q_1_*:*Q_2_*:*Q_3_* = 1:1:1 results in the corresponding layer thicknesses with ratio of *t_1_*:*t_2_*:*t_3_* ≈ 1.5:1:1.5 defined by the Poiseuille velocity profile. The flow streams flowing closer to the side walls occupy a larger portion of the channel width (37.5%), whereas the innermost stream flows faster with a smaller layer thickness (25%), despite having the same flow rate. A flat stacking of streams results from a non-inertial laminar flow with *Re* ≲ 1, whereas an inertial flow with *Re* ≈ *O*(20) will bend the stacked streams downwards due to the secondary Dean flow vortices (Figure S1 and Note S1). For a parallel inlet configuration (Σ*D_i_* = *W_c_*) with four inlets, a horizontal stacking of four flow streams is obtained with layer thicknesses, i.e., *t_1,4_* and *t_2,3_* equal 30.2% and 19.8% of *W_c_,* respectively, defined by the Poiseuille velocity profile such that *t_1,4_*:*t_2,3_* ≈ 1.5:1. An array configuration with four inlets, where two inlets are placed in parallel followed by another two in series, leads to a hybrid checkered flow profile with four uniformly distributed areas associated with each inlet flow.

To validate these numerically predicted flow profiles through experiment, we fabricated 3D HFL devices in which multi-layered flow streams are converged from mm-scale channel cross-sections into an order-of-magnitude smaller outlet channel (250 μm ྾ 250 μm) and structured microparticles are cured upon UV exposure through a photomask (Figures 2D and 2E). The microchannel aspect ratio, i.e., channel width *W_c_* to channel height *H_c_*, at the point of inlet entry can influence the flow profile within the square channel outlet, in addition to the inlet configuration (Figure S2). For consistency, we fixed the channel height, *H_c_* = 1.5 mm, for the experimental demonstrations unless otherwise mentioned, which results, for example, in *W_c_*:*H_c_* = 0.5, 2, and 1, respectively, for the inlet configuration (Σ*D_i_* = *W_c_*) depicted in Figures 2C and 2D. We have numerically investigated that additional variations of the sculpted flow profiles are possible for significantly higher channels, up to *H_c_* = 9 mm, and higher aspect ratios *W_c_*:*H_c_* = 3, combined with variable *W_c_*:*D_i_* = 0.5 to 4 (Note S2). However, these larger microchannel designs lead to increased delay time in a stop-flow cycle due to higher capacitance in the fluidic circuit. Therefore, we did not investigate these microchannel designs experimentally.

The shapes of fabricated microparticles match well with the numerically predicted cross-sectional flow profiles, owing to the meticulously crafted numerical models (Figure S3 and Note S3). For the first inlet configuration with Σ*D_i_* = *W_c_*, we cured multi-layered particles with vertical, horizontal, and hybrid flat layers stacking (Figure 2D). Beyond flat flow profiles, we also demonstrated a controlled creation of multi-layered upward or downward curved flow profiles by using the second and third inlet configurations (Figure 2E). We cured particle shapes resembling ‘*latte art*’ using symmetric and asymmetric inlet configurations for Σ*D_i_* < *W_c_* and Σ*D_i_* > *W_c_*. We have performed dimensional characterization to confirm high uniformity across particles and individual layers (Figure 2F). Moreover, the layer thicknesses measured experimentally from the fabricated particles matched nicely with the numerical prediction in Figure 2C, with <2% deviation. We estimated the pattern accuracy, i.e. the similarity between simulation predictions and experimental particle patterns, and the precision, i.e. the pattern uniformity between a batch of experimental particles, demonstrating the intersection-over-union (IoU) scores predominantly >90%, with a few cases at 86–89% attributable to diffusion-induced blurring at the layer interfaces and thresholding limitations (Figure 2G and Table S4).

#### 2.1.2 Flow sculpting using variable intra-channel pillar configurations

We also investigated the effect of pillar structures on flow sculpting within microfluidic channels (Figure 3). Full-height pillars, commonly fabricated via single-layer soft lithography in PDMS channels, have been used for inertial flow sculpting. However, their effectiveness emerges only at higher Reynolds numbers (*Re* > 10), where inertial effects break the fore-aft flow symmetry around cylindrical pillars, producing a net displacement of streamlines and enabling flow sculpting [31–36, 43, 44]. In contrast, non-inertial Stokes flows (*Re* < 1) remain largely unaffected by such pillars[30]. Partial-height herringbone structures have been used to mix or manipulate Stokes flows[42, 45–47], but their fabrication requires multi-step soft lithography with precise layer alignment, which is laborious. Our hybrid fabrication approach, combining 3D-printed molds with soft lithography, enables precise, flexible positioning of partial-height pillars, expanding the range of flow sculpting capabilities.

We examined how the symmetry of arbitrarily shaped pillars affects their ability to sculpt multilayered flow profiles (Figure 3A). A 2D shape (e.g., triangular) extruded along the Z-axis forms a full-height (*H_p_*:*H_c_* = 1) or partial-height (*H_p_*:*H_c_* < 1) pillar within the channel, where *H_p_* is the height of the pillar. At low Reynolds numbers (*Re* ≲ 1), both full- and partial-height symmetric pillars (about the YZ plane) do not significantly disturb such non-inertial flows with a checkerboard-shaped profile. However, at *Re* > 10, both pillar types generate inertial effects and Dean-like vortices that modify the incoming flow, consistent with inertial flow sculpting methods. In contrast, our approach operates in the Stokes regime (*Re* < 1), where flow manipulation relies on guiding co-flowing laminar streams using topological features that induce secondary flows in the YZ plane (Figure 3B). We show sculpted checkerboard-patterned flows attained via *advection maps*, which visualize the net displacement for a flow operator in the channel cross-sectional domain as 2D vectors. Symmetric pillars about the YZ plane cause reversible shaping of the flow^32, 44^, resulting in a net-zero sculpting effect due to the complementary deformation and relaxation of transverse displacement in the counterclockwise (CCW) and clockwise (CW) directions, respectively (Figure S4A and Note S4). To achieve irreversible flow sculpting in the Stokes regime, asymmetric structures about the YZ plane are necessary (Figure S4B). These include partial-height YZ-asymmetric pillars with symmetry or asymmetry about the XZ plane, which result in a pair of advection vortices or a single advection vortex (i.e., rotational displacements about some location in the YZ plane), respectively, as depicted in the advection maps (Figure 3B). Each oblique side of the triangular pillar (green or red) contributes to an advection vortex in a CW or CCW direction. By mirroring an XZ-symmetric triangular pillar about the YZ plane, i.e., from the triangle pointing downstream to the triangle pointing upstream, the direction of the advection vortex-pair pointing downwards is reversed to pointing upwards (Figure S4C). Similarly, the direction of the single CW advection vortex can be reversed to a CCW advection vortex by mirroring the XZ-asymmetric pillar about the YZ plane. Adjacent pillars can become hydrodynamically connected, generating an effective slope that induces an advection vortex, which persists until the pillars are sufficiently spaced apart (Figure 3C). Pillar features, such as depth (*H_p_*:*H_c_*), width (*W_p_*:*W_c_*), aspect ratio (*L_p_*:*W_p_*), edge bevel, and orientation, can be tuned to produce diverse sculpting effects (Figures S5 and Note S5). A composite channel design incorporating such partial-height (*H_p_*:*H_c_* < 1) pillars with different inlet configurations and superimposed CW/CCW vortices produces uniquely shaped structured microparticles (Figure 3D).

#### 2.1.3 Flow sculpting using variable outlet configurations

The geometry of the outlet channel, including its cross-sectional shape and dimensions, is crucial in defining the final cross-sectional outline of the produced microparticles and fibers, since the UV illumination is applied in this region to rapidly solidify the confined structured flow. Although existing techniques can fabricate microchannels with various cross-sectional boundary shapes[48, 49], they remain limited in achieving complex channel designs that integrate both geometrically defined outlets and multiple inlets. Our 3D-printed mold method effectively addresses this limitation: multi-inlet microchannel molds with diverse outlet geometries, e.g., inverted trapezoidal, triangular, slit-shaped, and square outlines, can be readily fabricated via 3D printing (Figure 4A). Moreover, the size of the flow profile, and consequently that of the resulting microparticles and fibers, can be tuned by adjusting the outlet channel dimensions.

Using this strategy, demonstrative structured microparticles of large (∼500 μm), medium (∼250 μm) and small (∼100 μm) sizes were fabricated and presented here (Figure 4B). The smallest particle size shown here is ∼100 μm. When the outlet channel size is further reduced, the primary challenge does not arise from the flow sculpting process, but rather from fabricating 3D-printed molds with dimensions below 100 μm, and from stopping the flow in such a small channel while maintaining the established flow profile. As the channel dimensions decrease, the hydraulic resistance increases, and the allowable operating window for stable flow stopping becomes narrower. However, smaller microparticles could be achieved by integrating with a sacrificed sheath flow in a 100 μm channel. As in any flow sculpting method, controlling the flow rate ratio remains the most straightforward means of adjusting the layer proportions within the flow profile (Figure 4C).

### 2.2 Representative fabrications of tailored microparticles and fibers

As one of the fundamental properties of microparticles or fibers, material composition plays a crucial role in defining their functionalities and thus guiding their potential applications. By integrating different material streams in 3D hydrodynamic flow lithography, a broad range of functional features can be achieved, e.g., solid or hollow interiors, functionalized surfaces, and localized expansion or contraction in response to specific stimuli, depending on the intrinsic material properties. For preliminary demonstrations, we fabricated several representative microparticles and fibers, with their material compositions and fabrication parameters summarized in Table S5. Firstly, we produced biocompatible PEGDA**-**GelMA particles with a three-layered pattern that can direct cell growth and intercellular interaction. As shown in time-lapse images, cells gradually migrated towards the particle and adhered exclusively to the GelMA regions[50], and finally binding the microparticles together (Figure 5A; see also Figure S10 and Movie S2). Secondly, we produced amphiphilic PEGDA**-**PPGDA particles with internal hydrophilic cavities that can facilitate the formation of uniform droplets without the need for conventional droplet microfluidics[6, 37, 38] (Figure 5B). Thirdly, we produced single-material PEGDA particles and fibers with structured cross sections and anisotropy along the fiber length, which is incorporated by tuning the UV exposure through the photomask (Figure 5C). Lastly, we realized a thermally-responsive PEGDA**-**PNIPAM bilayer fiber that exhibited reversible shrinkage and curling behavior as the temperature was increased from room temperature to 60 °C[51] and back (Figure 5D). With these representative examples of microparticles and fibers, we aim to demonstrate the platform’s compatibility with commonly used photocurable materials, as well as its capability to fabricate both fibers and particles with complex structures.

### 2.3 Fast flow profile engineering via software

All numerical work shown thus far was conducted using full 3D CFD simulations in COMSOL, which requires a laborious sequence of 3D channel design, meshing, simulation, and post-processing (see steps 1 and 2 in section 3.1). Prior work has shown that similar flow sculpting modalities can be encoded into lightweight computational models using only 2D advection maps [52, 53] or machine learning-based prediction [56], and we sought to apply a similar methodology to 3D HFL to reduce the time needed for numerical exploration and design.

Three features of 3D HFL complicate direct adoption of these approaches. First, prior works hold channel aspect ratio fixed, enabling matched domain sizes across operators, which is useful for chaining advection maps and a convenience for machine learning models, whereas 3D HFL uses channel aspect ratio variation as part of its sculpting toolkit and to span a range of manufacturing lengthscales. More critically, prior work assumes fixed-inflow designs, in which a single inlet configuration (which could comprise multiple streams) is shaped by independent downstream operators. 3D HFL instead uses inlets as mid-sequence operators, making them sequence-position-dependent: the flow transformation produced by a new inlet depends on how many streams already occupy the channel at that point, including both the initial stream count and any streams added by preceding inlets. Finally, the full 3D HFL operator space - single- and double-slope pillar variants with varying size, shape and depth; operators placed in expanded or reduced channels; and composite multi-pillar configurations (see Figure 3) - is vast, difficult to enumerate, and non-trivial to curate exhaustively into a single precomputed library.

The aspect ratio challenge can be addressed by introducing transitional advection maps that rescale flow profiles between operators of different channel shape and dimension, and sequence dependence can be handled by constructing library entries that explicitly encode the stream count at the point of inlet insertion. To test these two solutions, we developed a forward model, graphical user interface (GUI, Figure 6), and scoped advection map library using the same advection map approach described in Ref [53]. This scoped library covers single-slope pillars, inlet operators, and five outlet shape transitions (square, slit, trapezoidal, and two triangles), with each advection map generated using the same CFD model described in section 3.1. The total operator count in one sequence is capped at 5, encompassing both pillars and inlets, to keep the maximum number of simultaneous inlet streams experimentally practical. Each pillar is parameterized by 7 values (channel width, channel height, pillar depth, and 4 coordinates describing the pillar’s vertex locations), yielding 900 distinct pillar configurations, and 900^5^∼10^14^ possible combinations for a full 5-pillar sequence. We validated this forward model against the CFD model, demonstrating accurate prediction capability with substantially reduced computation time (Figure 6); the representative GUI design parameters used to generate the flow profiles in Figure 4D are provided in Figure S9.

**Figure 6.**
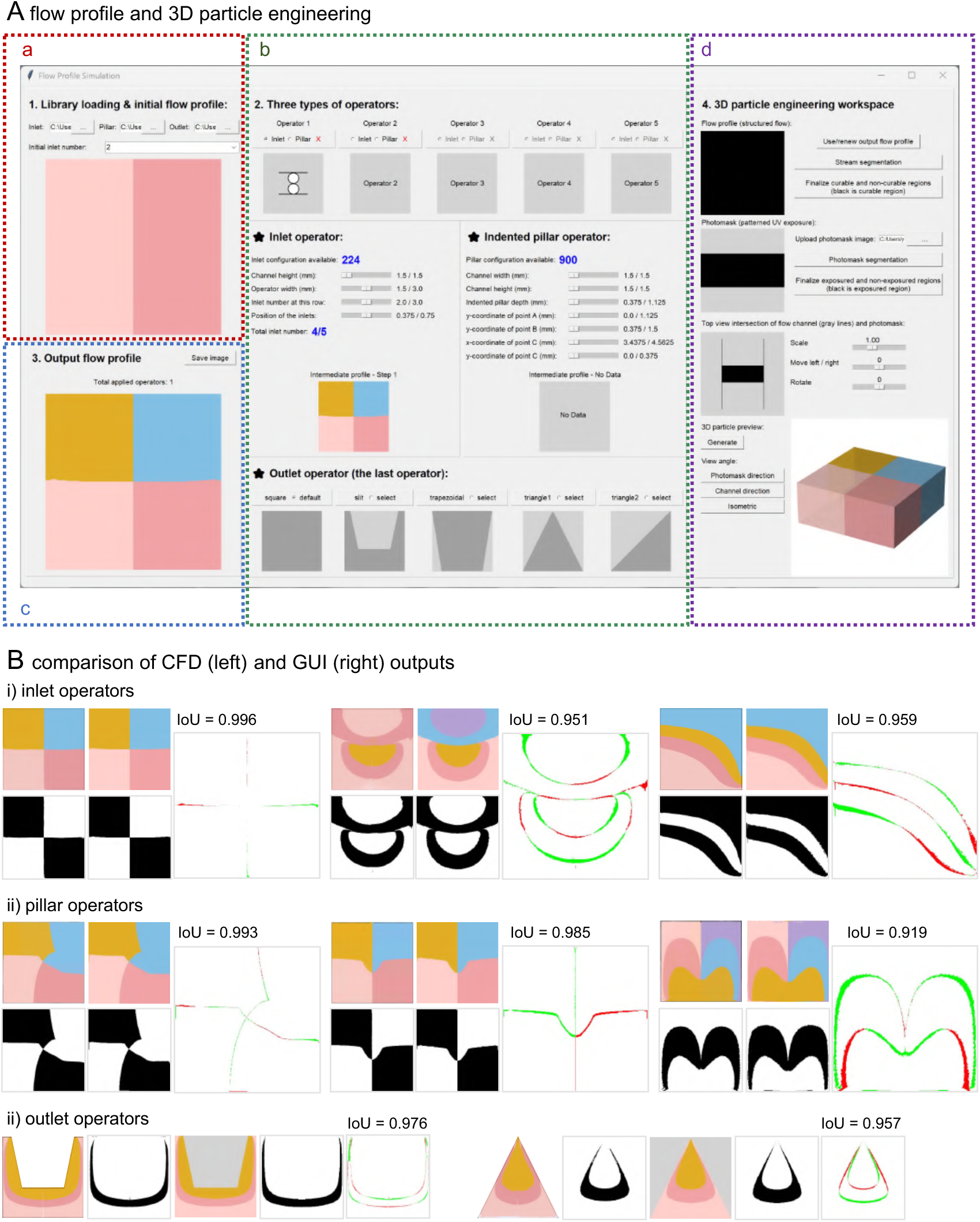
Graphical User Interface (GUI) for 3D hydrodynamic flow sculpting. (A) The GUI consists of four panels. Panel (a) allows users to select an initial flow profile containing up to 5 horizontally stacked streams (inlets in parallel). Panel (b) enables further vertical stream stacking through additional inlet configurations (inlets in series), flow sculpting via indented triangular pillar configurations, and geometry constraining through outlet configurations. These configurations can be applied either independently or in combination within the panel. Intermediate flow profiles generated at each operation step are displayed accordingly. Panel (c) presents the resulting flow profile after all operations. Panel (d) provides a 3D visualization function by intersecting the engineered multilayered flow pattern with a patterned photomask for 3D microparticle engineering. (B) Comparison between CFD-predicted (left in each pair) and GUI-predicted (right in each pair) flow profiles for inlet, pillar, and outlet operators. Corresponding binary masks, contour overlaps (red indicates regions present only in the CFD mask, while green indicates regions present only in the GUI mask), and IoU scores are shown to quantify prediction accuracy.

Expanding this library to cover double-slope pillars, intermediate channel shape transitions, and composite configurations would require significant additional simulation and careful parameterization of a far more complex design space. These are tractable but resource-intensive tasks, and the single-slope library demonstrated here is sufficient to establish the viability of the forward modeling approach and validate its predictions against CFD-simulated flow profiles (Figure 6); these CFD predictions were, in turn, validated against experimental results (Table S4). In its current form, the GUI supports the following operator classes: single-slope partial-height pillar operators, inlet operators (series and parallel configurations), and five outlet shape transitions (square, slit, trapezoidal, and two triangle variants); double-slope pillars, intermediate channel shape transitions, and composite multi-pillar configurations are not yet included in the library. This scope limitation should be considered when interpreting the forward-prediction results shown in Figure 6.

## 3. Materials and Methods

### 3.1 Workflow

The overall flow profile design is guided by CFD simulations, which predict the cross-sectional flow profiles created by microchannel design with different inlet, pillar, and outlet configurations. These simulations provide the primary framework for determining the flow sculpting strategy (Figure 7). While the device assembly, precursor compositions, and operational parameters, such as flow rates and curing conditions, are experimentally optimized to ensure stable operation and reliable particle fabrication.

**Figure 7.**
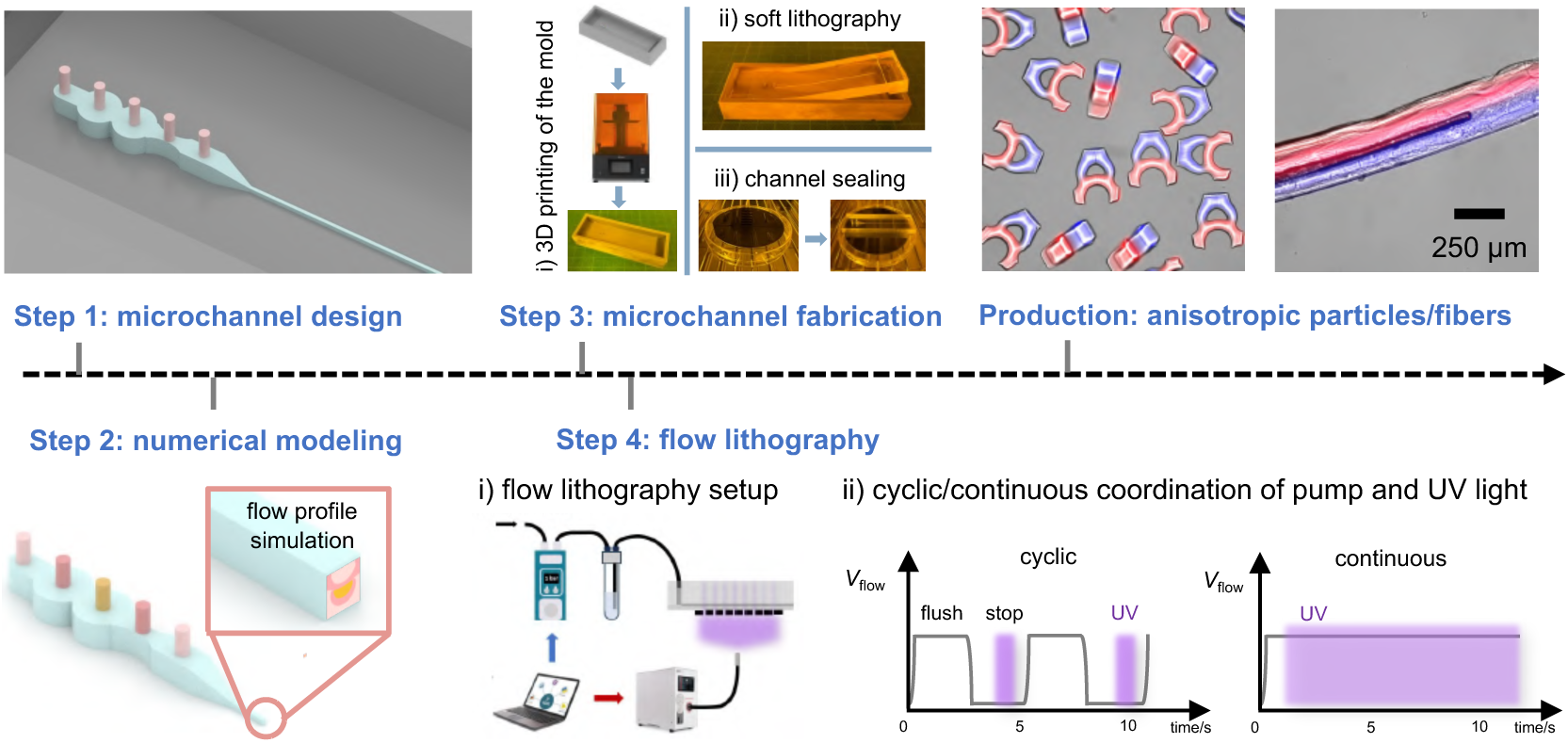
Workflow for 3D hydrodynamic flow lithography.

#### 3.1.1 Step 1: microchannel design

The microchannel model was designed using 3ds Max^®^ computer-aided drafting software (Autodesk Inc.) and exported as the SAT file for numerical simulation or the STL file for 3D printing. The following general dimensions and guidelines were used for the microchannel design:

1. **Inlet pillars:** The inlet diameter was typically set as 0.75 or 1.5 mm to match the outer diameter of the inlet tubing. For designs with parallel inlets, the inlet pillars were extended beyond the mold wall height with a smaller top (0.6 mm) to eliminate the need for hole punching in the PDMS replica and prevent liquid leakage at the inlet junctions.
2. **Main channel body:** The main channel width was set as 0.75, 1.5, or 3 mm based on the inlet diameter, number, and configuration, with a fixed channel height of 1.5 mm.
3. **Transition section:** A gradually converging section, with a length of 2.5 mm and an original height of 1.5 mm, was incorporated to connect the main channel and the outlet channel, ensuring a smooth hydrodynamic flow transition.
4. **Outlet channel:** The outlet channel length was typically set as 2 mm in the numerical model and 35 mm for the 3D printing model.

#### 3.1.2 Step 2: numerical modeling

The SAT file containing the microchannel design was imported into COMSOL^®^ Multiphysics for numerical analysis. The Single-Phase Laminar Flow module was used to predict the flow profile and estimate pressure and velocity distributions. All simulations were performed using a working fluid composed of 60 vol.% PEGDA and 40 vol.% ethanol (density: 987 kg·m^−3^; viscosity: 7.55 mPa.s). The colors and positions of the inlet pillars in the numerical model correspond to those of the streamlines, enabling clear visualization of the layered flow profile. The inlet boundary condition was usually set as a flow rate of 5.7e-10 m^3^/s per inlet for channels with a 250 µm × 250 µm square outlet, unless otherwise specified. Additional pillars (diameter: 0.05 mm, length: 3.54 mm) were added to mimic inlet tubings with an inner diameter of 0.18 mm and a length of 60 cm in the high-flow-resistance model (Figure S3C). The Reynolds number is defined as *Re* = ρ*UW*_c_/μ, where ρ is the fluid density, *U* is the mean velocity at the outlet channel, *W*_c_ is the outlet channel width (characteristic length), and μ is the dynamic viscosity. For the standard operating conditions used here (flow rate 5.7 × 10^−10^ m^3^/s per inlet, 250 µm outlet), *Re* ≲ 1, confirming operation in the non-inertial Stokes flow regime throughout this work.

#### 3.1.3 Step 3: microchannel fabrication

1. **3D printing of the microchannel mold:** After obtaining a satisfactory flow profile from the numerical simulation, the microchannel design was finalized by adding mold walls and adjusting the outlet channel dimensions. The finalized mold design was exported as an STL file and converted to sliced CTB files using CHITUBOX^®^ software. The CTB file was then imported to the 3D printer (Sonic MINI 8K and Aqua-Gray 8K Resin, PHROZEN TECH CO., LTD) for microchannel mold printing. For the 500 μm × 500 μm square outlet channel, the design specified a channel width of 500 μm and a height of 530 μm to compensate for the offset in the Y-axis of the 3D printing. The printing parameters were set to a printing layer thickness of 50 μm and an exposure time of 2.1 s per layer. For the 250 μm × 250 μm outlet channel, the dimensions were adjusted to 250 μm × 325 μm, with a layer thickness of 25 μm and 1.0 s exposure time. For the 100 μm × 100 μm outlet channel, the design dimensions were 100 μm × 160 μm, printed with a layer thickness of 20 μm and 1.4 s exposure time.
2. **Surface treatment of the microchannel mold:** The 3D-printed mold was then thoroughly washed with isopropanol, dried with an air flush, and subjected to a UV post-curing for 60 minutes (Anycubic Wash & Cure 2.0). After a 1-minute plasma activation, the mold was exposed to the vapor atmosphere of 50 μL Trichlor(1H,1H,2H,2H-perfluorooctyl)silane (448931, Sigma-Aldrich) for 20 minutes in a Petri dish, to complete the hydrophobic surface treatment.
3. **Microchannel fabrication:** The PDMS microchannel was fabricated using soft lithography. Briefly, the PDMS base and curing agent (184 Silicone Elastomer Kit, Dow Inc.) were mixed at a 5:1 mass ratio and stirred until homogeneous. Then the mixture was poured over the microchannel mold, degassed, and cured at 65°C for 1 h (90 °C was used for the first curing step to clean the mold surface). After curing, the PDMS replica was removed from the mold, cut off both step ends, and the inlet holes were perfectly extended by vertical punching.

For channel sealing, a 100-150 μm PDMS thin membrane was prepared by spin coating the PDMS mixture onto a silicon wafer at 500 rpm for 30 s, followed by curing at 65 °C for 15 min. The PDMS channel replica and thin membrane were then bonded using oxygen plasma treatment for 1 min, forming a sealed PDMS microchannel.

After detaching the sealed channel from the silicon wafer, an additional cover glass (20 × 26 × 0.4 mm; Paul Marienfeld GmbH & Co. KG) was attached beneath the PDMS thin membrane at the inlet and converging section (excluding the outlet channel) using a very thin layer of uncured PDMS mixture as an adhesive. The assembled device was further cured at 90 °C for 24 h to enhance rigidity. Alternatively, a thin cover glass pre-cured with a thin PDMS membrane can also be used to seal the entire channel, though the additional glass layer at the outlet region would slightly affect the UV exposure. Finally, an open outlet for particle collection was created by cutting at the end of the outlet channel.

#### 3.1.4 Step 4: flow lithography

The FL setup consists of a laptop, a PDMS microchannel chip connected with PEEK inlet tubings, a fluid driving system (LineUp Flow EZ 7000, Fluigent Deutschland GmbH), and a UV curing unit (S2000 elite, Excelitas Canada Inc.). The precursor flow is first introduced and shaped at the sculpting channel and subsequently solidified as microparticles or fibers at the outlet region under the UV exposure. A custom Python script was developed to automate the system, synchronizing the signals sent to both the UV light source and the pressure controllers. Specifically, SFL achieves particle fabrication through the cyclic coordination of flow stoppage and UV exposure[6, 37, 38, 54], whereas CFL enables fiber fabrication by keeping the flow continuous under UV exposure[41]. The UV light source was coupled with a collimator at an adjusted working distance to ensure a uniform exposure on the precursor flow across the entire photomask area. 100% UV light intensity was used in all experiments to maintain high UV exposure uniformity. A key consideration in the SFL configuration is the management of the high hydraulic flow resistance within the microfluidic system. As the flow was regulated using the pressure controller without additional mechanical valves, the hydrostatic pressure from the height difference between the pressure controllers’ reservoirs and the microfluidic chip can induce unwanted flow continuation even when the applied pressure is removed, which is detrimental to the SFL process [54]. So, tubings with different I.D.s (PEEK tubings with 1/32-inch (multi-inlet channel) and 1/16-inch (single-inlet channel) O.D., and 250 μm, 180 μm, and 130 μm I.D., Analytics-Shop) were intentionally chosen based on the size of the microchannel (Table S2) to appropriately increase flow resistance so as to minimize the unwanted flow and sustain uniform particle production for a long time. The photomask was attached directly underneath the PDMS thin membrane at the outlet channel area for particle fabrication. Any bubbles trapped inside the channel are detrimental to flow sculpting, so before connecting the tubing with the channel, the air inside the channel is expelled and the channel filled with ethanol or water. The environmental temperature for the GelMA particle fabrication was controlled at around ∼ 30 °C. For continuous fabrication of long fibers, there is no slit pattern in the photomask mask and except for the region of the outlet channel, the rest of the chip is covered to control the exposure dosage.

#### 3.1.5 Particles/fibers collection

The produced particles/fibers were collected from the outlet channel, thoroughly washed three times with ethanol to remove uncured precursor and water insoluble photoinitiator, and then stored in either ethanol or a 0.1% Pluronic^®^ F-127 (P2443, Sigma-Aldrich) aqueous solution to prevent aggregation or adhesion on the surface of the container. The particles for the cell experiment were washed and stored in cell culture medium.

### 3.2 Precursor solution preparation

The precursor solution consisted of 60 vol.% Poly (ethylene glycol) diacrylate (PEGDA, *M_w_*575, 437441, Sigma-Aldrich) and 40 vol.% ethanol (1.00983, Sigma-Aldrich), additionally with 5 vol.% photoinitiator (2-hydroxy-2 methylpropiophenone, Darocur 1173, 405655, Sigma-Aldrich) and 50 µg/mL fluorescent dyes (red: Methacryloxyethyl thiocarbamoyl rhodamine B, 23591, Polysciences, Inc. Headquarters; blue: Methacrylate-modified blue-emitting dye, MA-P4VB: 2-(methacryloyloxy)propyl 6-(4-methoxy-2,5-bis((E)−2-(pyridin-4-yl)vinyl)phenoxy)hexanoate)), provided by a collaborating laboratory), or 5 vol.% fluorescent nanoparticles dispersion solution, to visualize the particle structures (Table S3). For multiple-material particles, the compositions of the precursor solution are listed in Table S5, including polyethylene glycol (PEG, *M_w_* 200, P3015, Sigma Aldrich), polypropylene glycol diacrylate (PPGDA, *M_w_* 800, 455024, Sigma-Aldrich), Gelatine-Methacryloyl (GelMA, 900622, Sigma-Aldrich), N-Isopropylacrylamide (NIPAAM, H66262.22, Thermo scientific). GelMA precursor was prepared in the ∼ 50 °C water bath to melt and homogenize the solution, under the cover of aluminum foil. Density matching among different precursor streams is critical for maintaining stable flow patterns. A density mismatch between streams can lead to noticeable deviations between the simulated and experimentally obtained structures. To ensure flow stability and fidelity to theoretical designs, the densities of the precursor solutions should be carefully adjusted within ∼1% during preparation (PEGDA: 1.12 g/mL, PPGDA: 1.01 g/mL, PEG: 1.127 g/mL,ethanol: 0.79 g/mL, 60 vol.% PEGDA & 40 vol.% ethanol: 0.987 g/mL).

### 3.3 Cell culture

Human embryonic kidney cell line HEK293T was obtained from the American Type Culture Collection (ATCC, Manassas, VA, USA). The GFP-expressing U87 glioblastoma cell line was kindly provided by a collaborating laboratory. U87 and HEK293T cells were cultured in high-glucose Dulbecco’s Modified Eagle Medium (DMEM, GlutaMAX™ Supplement; Gibco™, Thermo Fisher Scientific, Waltham, MA, USA) supplemented with 10% (v/v) heat-inactivated fetal bovine serum (FBS; Sigma-Aldrich, Merck KGaA, Darmstadt, Germany) and 1% (v/v) penicillin–streptomycin (10,000 IU/mL penicillin and 10 mg/mL streptomycin; Sigma-Aldrich). All cells were maintained at 37 °C in a humidified atmosphere containing 5% CO₂ and were routinely tested for mycoplasma contamination.

For passaging, the culture medium was aspirated, and the adherent cells were washed once with phosphate-buffered saline (PBS). Cells were then detached by adding 5 mL of trypsin–EDTA solution (Gibco™) and incubating at 37 °C for 2–5 minutes, depending on the cell line, until detachment was observed. The enzymatic reaction was stopped by adding an equal volume of complete medium. The cell suspension was centrifuged, resuspended in fresh culture medium, and seeded into new culture flasks for further growth.

### 3.4 Cell-particle experiment protocol and imaging

Particles, in quantities ranging from 50 to 80, were seeded into non-treated 48-well plates containing cell culture medium. For each cell type, two wells were prepared as experimental samples, while a third well served as a negative control. A volume of 100 µL of cell suspension (containing 2 × 10⁶ cells/mL) was added to each well.

The plate was maintained under environmental control conditions (37 °C, 5% CO₂) on a Leica LAS X fluorescence microscope. Time-lapse imaging was performed over a period of 72 hours, with Z-stack images captured at 3-hour intervals to monitor cell behavior and interactions.

### 3.5 Statistical Analysis

Microparticle dimensions (width, height, and thickness) were measured from the microscopy images of particles dispersed in ethanol or water. For each sample, 15 individual particles were measured and reported as mean ± standard deviation (SD) (Table S4, Figure S6, and Figure S7). Microchannel dimensions were measured from microscopy images of the microchannel cross-sections. For each sample, 15 individual cross-sections were measured and reported as mean ± standard deviation (SD) (Figure S7).

The Flow pattern accuracy, the agreement between simulated and experimental particle cross-sectional patterns, was quantified using the Intersection-over-Union (IoU). One representative particle image per pattern was randomly selected and compared against the corresponding CFD prediction (Figure 2G, Figure S8, and Table S4). The use of n = 1 for accuracy measurement is justified because the CFD prediction is deterministic: for a given channel geometry and flow-rate ratio, the simulated cross-sectional flow profile is unique, so a single experimental measurement is sufficient to assess agreement with the prediction; particle-to-particle variability is captured separately by the precision metric.

For the Flow pattern precision, the agreement within a batch of microparticles, n ≥ 10 particles per batch, was used to compute pairwise IoU values, reported as mean ± SD (Figure 2G, Figure S8, and Table S4). The image processing and IoU computation were conducted using Python code. The results are reported in Table S4.

## 4. Conclusions

In conclusion, 3D hydrodynamic flow lithography shows great potential in the design and continuous fabrication of multi-material, anisotropic microparticles and fibers. We have demonstrated 3D HFL as a versatile platform capable of: 1) precise flow profile sculpting of up to five co-flowing streams with tunable sizes (∼100–500 μm), diverse outlines, and complex internal structures using variable inlet, pillar, and outlet configurations; 2) simplified, cleanroom-free microdevice manufacturing using low-cost 3D-printed molds to produce PDMS channels with precise topologies; 3) inertia-free and sheath-flow-free operation, ensuring stable performance across various flow velocities while improving material utilization; 4) broad material compatibility (e.g., PEGDA, PNIPAM, GelMA, and PPGDA) for fabricating functional, application-tailored microstructures; and 5) rapid, GUI-assisted forward modeling based on advection-map computation to predict flow profiles with substantially reduced computational cost. Together, these advancements establish 3D hydrodynamic flow lithography as a versatile and high-throughput platform—exceeding 10^4^ particles/hour via stop flow lithography—for engineering anisotropic microparticles and fibers with customizable size, structure, and composition. This throughput figure applies to the SFL particles demonstrated here and compares favourably with existing z-structuring methods such as lock-release, vertical, and soft-membrane flow lithography, which typically operate at *O*(10^2^) particles/hour; the two-orders-of-magnitude advantage arises primarily from the sheath-free, single-layer PDMS channel design that shortens each stop-flow cycle.

Nevertheless, several opportunities for further extension remain. Channel geometries that expand from bottom to top, exhibit deep concave features, or place structures at the bottom of the channel are inherently restricted by the current PDMS-on-mould fabrication workflow. Direct 3D printing of oxygen-permeable channels could potentially overcome these geometric limitations and broaden the design possibilities in 3D hydrodynamic flow lithography. Moreover, the resolution of the 3D-printed molds or microchannels has a significant influence on the sculpted flow structures. Building on the forward-prediction GUI introduced here, a natural extension would be a method for inverse-design that automatically determines 3D-HFL device parameters that produce a target flow pattern, possibly using standard optimization techniques (e.g., the genetic algorithm [56, 57]) or artificial intelligence-based approaches [55].

## Supplementary Material

See the supplementary material for additional supporting movies, figures, and notes.

## Acknowledgment

Y.Z. gratefully acknowledges the Chinese Scholarship Council for the doctoral fellowship and the support from all members working in the Control and Manipulation of Microscale Living Objects group and the Heinz Nixdorf Chair of Biomedical Electronics. We also thank Dr. Sergei I. Vagin for kindly providing the blue-emitting dye and thank Dr. Morteza Kafshgari for kindly providing different cell lines.

## Data Availability Statement

The data that support the findings of this study are available from the corresponding author upon reasonable request. The GUI and the library data underlying the predictive design framework (Figure 6, Movie S1) are openly available at https://github.com/yiyingzou/3D-hydrodynamic-flow-sculpting.

## Author Contributions

Y.Z. designed and performed the experiments, carried out the CFD simulations, analysed the data, and wrote the manuscript. M.A.S., M.Z.U.K., and M.U.A. contributed to the device fabrication, flow lithography setup, and characterisation. D.S. contributed to the advection-map framework and the GUI design. G.D. conceived and supervised the project, secured funding, and edited the manuscript. All authors have read and approved the final version of the manuscript.

---

**Table S1.** Summary of droplet microfluidics and representative SFL approaches for anisotropic microparticle fabrication.

| Approach | Structure control method | Contribution in structure creation | Channel fabrication | Multi-material | Throughput /particles·h<sup>-1</sup> | Particle size range /μm | Ref. |
| --- | --- | --- | --- | --- | --- | --- | --- |
| Droplet microfluidics | Interfacial morphology | 3D interface-derived shapes | Polymer base, glass capillary based, needle based, etc. | Yes | 10<sup>2</sup> – 10<sup>3</sup> Hz | cover the range from the nano to micro scale | [1] |
| Stop-flow lithography | Photomask; elementary flow profile | Structure created at X-Y plane by UV pattern and at Y-Z plane by flow profile | Photolithographic mold; single-layered PDMS channel | Yes | 10<sup>7</sup> | *O*(1)-*O*(100) | [2] |
| Two-step lithography | Twice exposures with digital micromirror device | Structures assembled at X-Y plane and along Z direction |  | Yes | / | *O*(100) | [3] |
| Lock release lithography | Interior structure of the channel | Anisotropy in Z direction |  | Yes | 10<sup>3</sup> | *O*(100) | [4] |
| Interference lithography | 2D phase mask; amount of inhibitor and UV light dose |  |  | Yes | / | *O*(10) | [5] |
| Non-uniform UV lithography | Non-uniform UV absorption with UV absorber in different concentrations |  |  | Yes | / | *O*(100) | [6] |
|  |  |  |  | No | / | *O*(10) | [7] |
|  | Non-uniform UV exposure with different focus planes |  |  | No | / | *O*(10)-*O*(100) | [8] |
| Soft membrane lithography | 3D layer by layer dynamic photomask projection with a digital micromirror device | 3D structure | Photolithographic mold; two-layered PDMS channel | Yes | / | *O*(100) | [9] |
| Vertical flow lithography |  |  | 3D printed mold; 3D PDMS channel |  | 10<sup>2</sup> | *O*(10)-*O*(100) | [10] |

| Approach | Structure control method | Contribution<br>in structure creation | Channel fabrication | Multi-material | Throughput<br>/particles·h<sup>-1</sup> | Particle size<br>range /μm | Ref. |
| --- | --- | --- | --- | --- | --- | --- | --- |
| Hydrodynamic<br>flow lithography | Stacking multiple streams<br>from different positions or<br>direction | Structure created at Y-Z plane by advanced<br>flow profile sculpting | Photolithographic mold;<br>two-layered PDMS channel | Yes | / | *O*(10)-*O*(100) | [11] |
| Inertial flow<br>lithography | Sculpting the inertial flow<br>using sequenced obstacle<br>pillars inside microchannel |  | Photolithographic or<br>stereolithographic mold;<br>single-layered PDMS<br>channel | Yes | 10 | *O*(100)-<br>*O*(1000) | [12,<br>13] |
|  |  |  |  |  | / | *O*(100) | [14] |
|  |  |  |  |  | 10<sup>2</sup> | *O*(100)-<br>*O*(1000) | [15] |
|  |  |  |  |  | 10<sup>4</sup> | *O*(100) | [16] |
|  |  |  |  |  | 10<sup>4</sup> | *O*(100) | [17] |
| Co- flow<br>lithography | Sculpting the non-inertial<br>flow using 3D printed inlet<br>connector |  | 3D printed inlet connected<br>to a glass capillary or tubing | Yes | 10<sup>4</sup> | *O*(100) | [18-20] |
| Structured flow<br>lithography | Sculpting the outline of the<br>flow using microchannel<br>with specific boundary<br>shape |  | COC with aluminum rod<br>template inside; thermal<br>fiber drawing of COC<br>channel until to the target<br>dimensions | No | 10<sup>4</sup> | *O*(10)-*O*(100) | [21] |
|  |  |  | Convex 440C stainless steel<br>metal molds; PDMS soft<br>lithography and channel<br>assemble | No | 10<sup>3</sup> | *O*(100) | [22] |

**Table S2.**
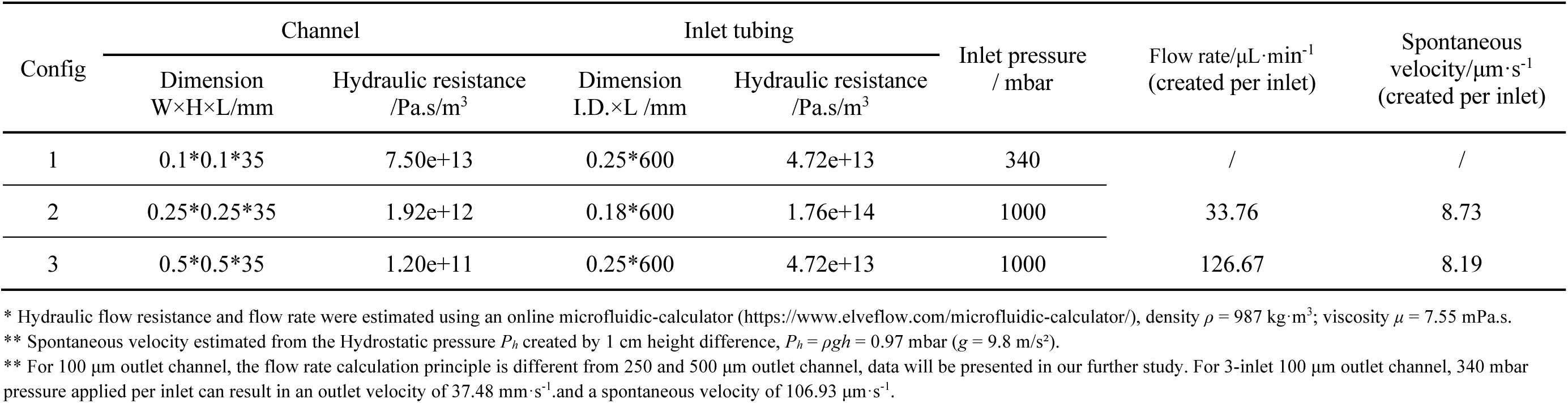
Microchannel and inlet tubing configurations.

**Table S3.**
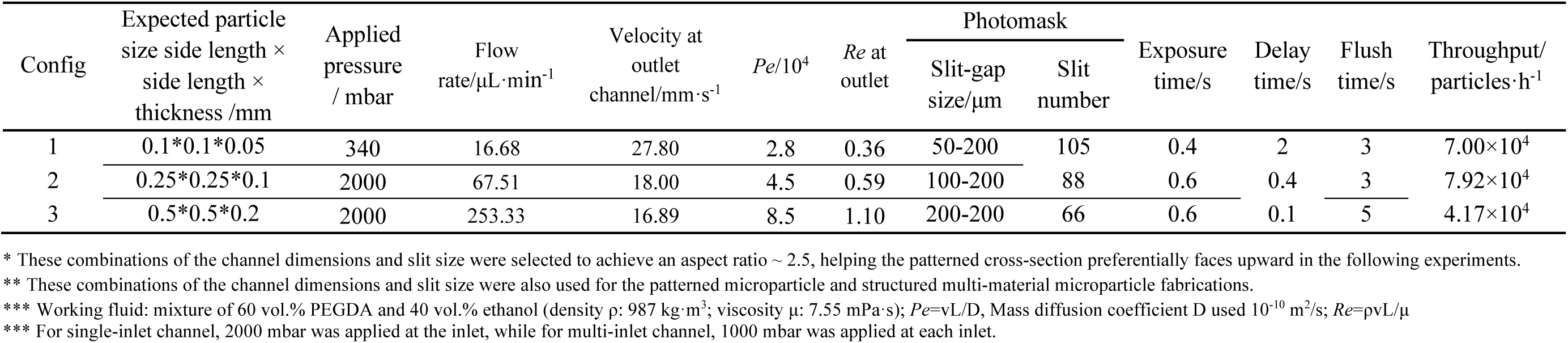
Parameters for blank microparticles fabricated with single-inlet configurations.

| Config | Expected particle<br>size side length ×<br>side length ×<br>thickness /mm | Applied<br>pressure<br>/ mbar | Flow<br>rate/$\mu\text{L}\cdot\text{min}^{-1}$ | Velocity at<br>outlet<br>channel/$\text{mm}\cdot\text{s}^{-1}$ | $Pe/10^4$ | $Re$ at<br>outlet | Photomask | | Exposure<br>time/s | Delay<br>time/s | Flush<br>time/s | Throughput/<br>particles·h<sup>-1</sup> |
| --- | --- | --- | --- | --- | --- | --- | --- | --- | --- | --- | --- | --- |
| | | | | | | | Slit-gap<br>size/$\mu\text{m}$ | Slit<br>number | | | | |
| 1 | 0.1*0.1*0.05 | 340 | 16.68 | 27.80 | 2.8 | 0.36 | 50-200 | 105 | 0.4 | 2 | 3 | $7.00\times 10^4$ |
| 2 | 0.25*0.25*0.1 | 2000 | 67.51 | 18.00 | 4.5 | 0.59 | 100-200 | 88 | 0.6 | 0.4 | 3 | $7.92\times 10^4$ |
| 3 | 0.5*0.5*0.2 | 2000 | 253.33 | 16.89 | 8.5 | 1.10 | 200-200 | 66 | 0.6 | 0.1 | 5 | $4.17\times 10^4$ |

\* These combinations of the channel dimensions and slit size were selected to achieve an aspect ratio $\sim 2.5$ , helping the patterned cross-section preferentially faces upward in the following experiments.
\*\* These combinations of the channel dimensions and slit size were also used for the patterned microparticle and structured multi-material microparticle fabrications.
\*\*\* Working fluid: mixture of 60 vol.% PEGDA and 40 vol.% ethanol (density $\rho$ : $987 \text{ kg}\cdot\text{m}^{-3}$ ; viscosity $\mu$ : $7.55 \text{ mPa}\cdot\text{s}$ ); $Pe=vL/D$ , Mass diffusion coefficient $D$ used $10^{-10} \text{ m}^2/\text{s}$ ; $Re=pvL/\mu$
\*\*\* For single-inlet channel, 2000 mbar was applied at the inlet, while for multi-inlet channel, 1000 mbar was applied at each inlet.

**Table S4.**
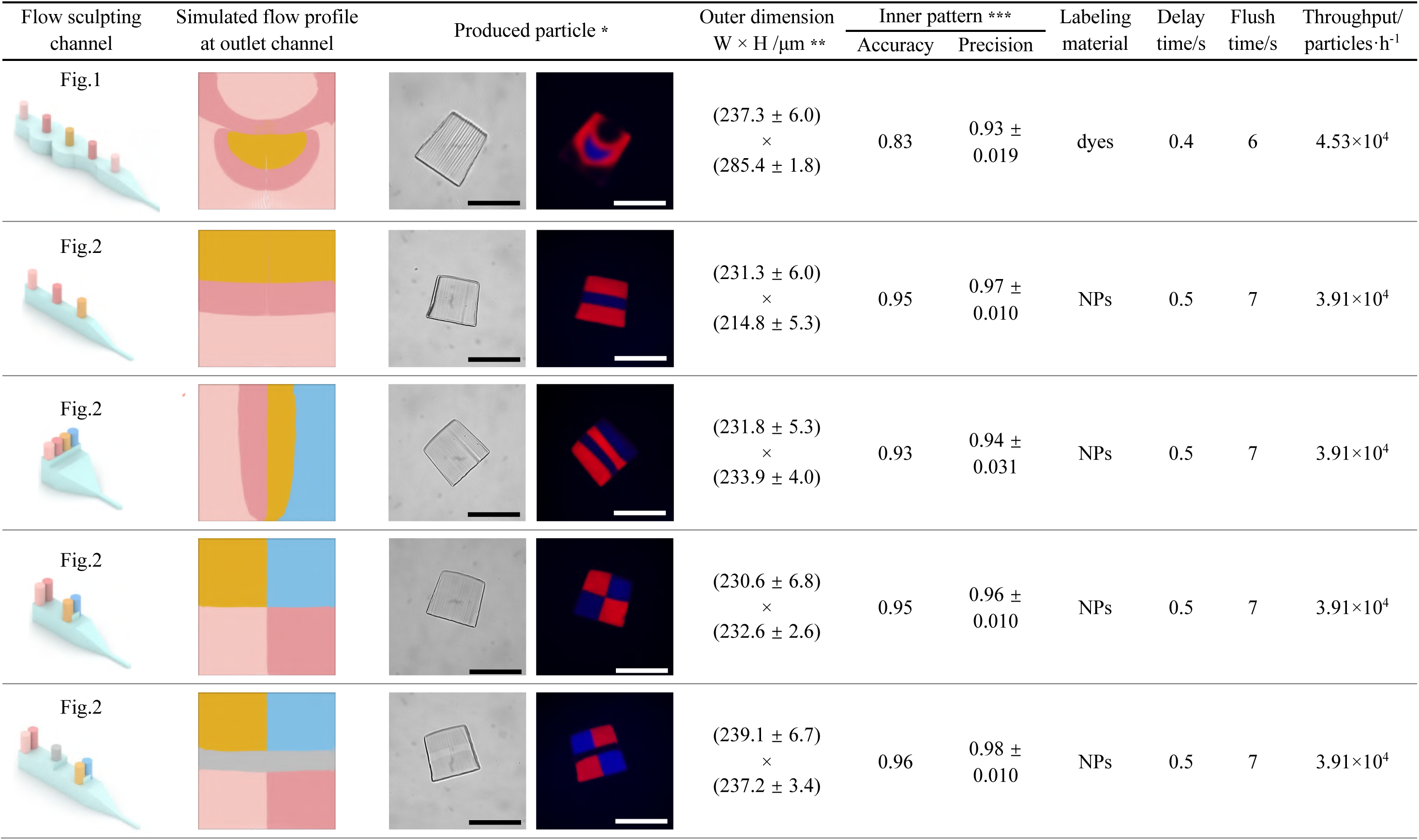
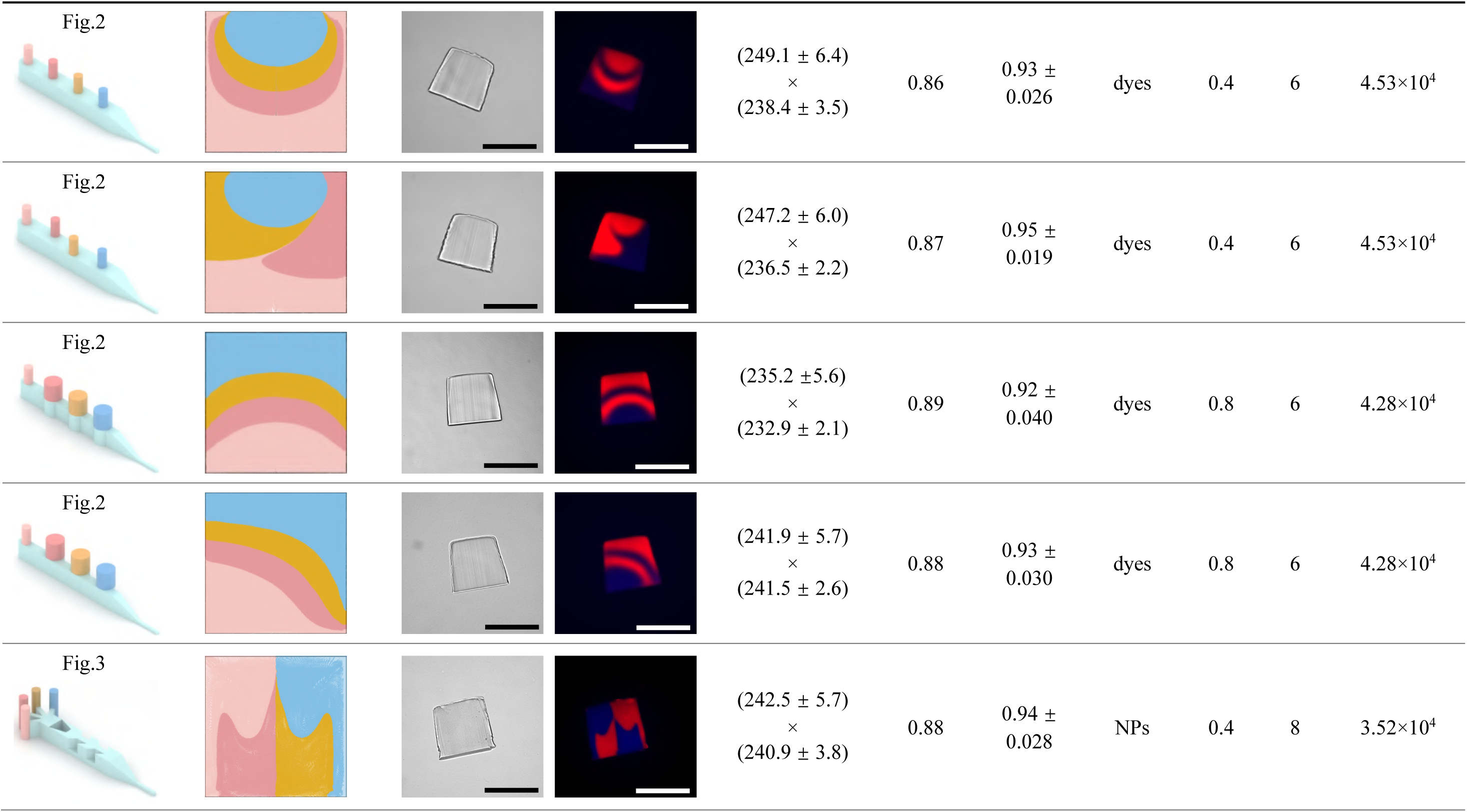
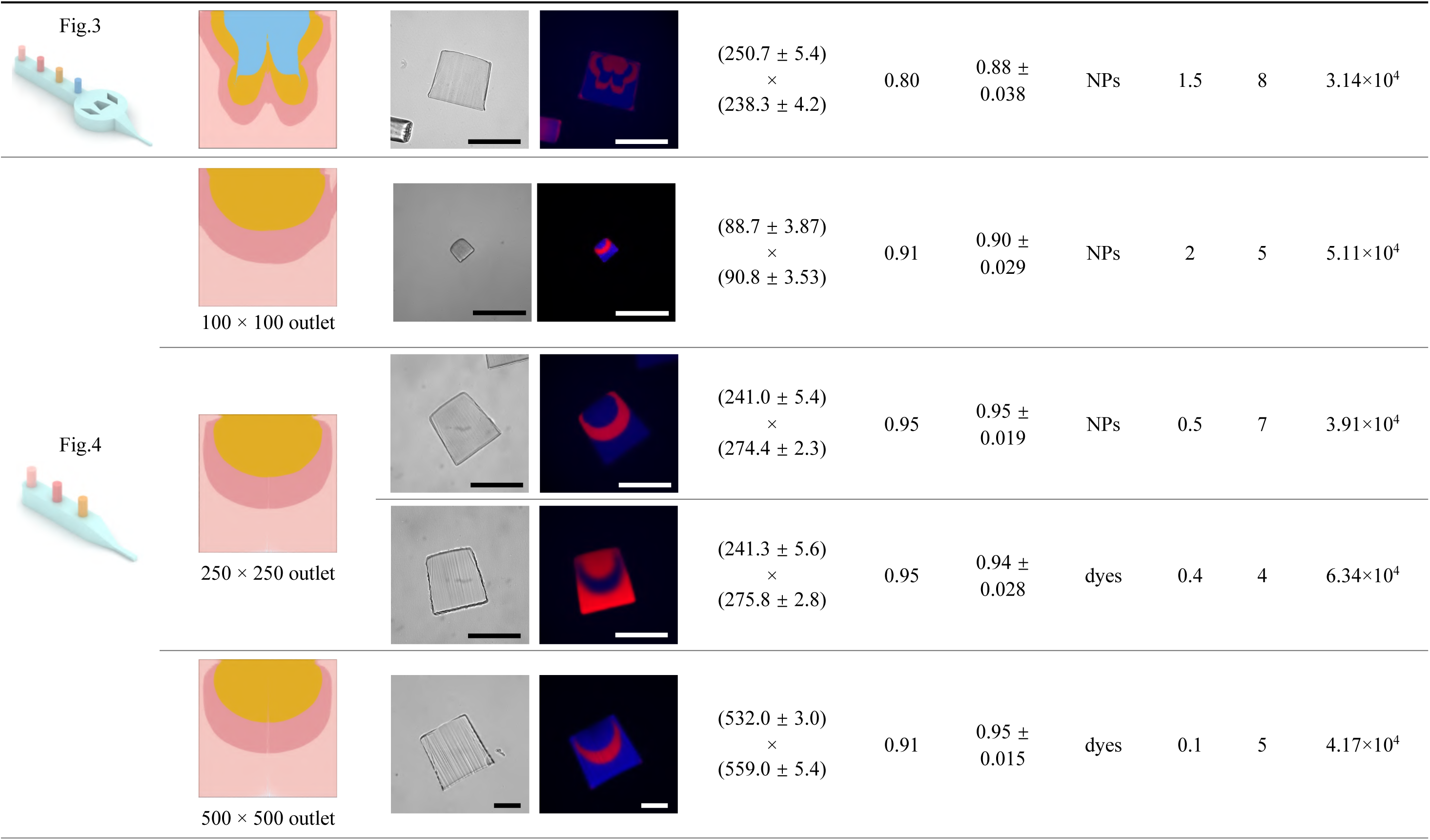
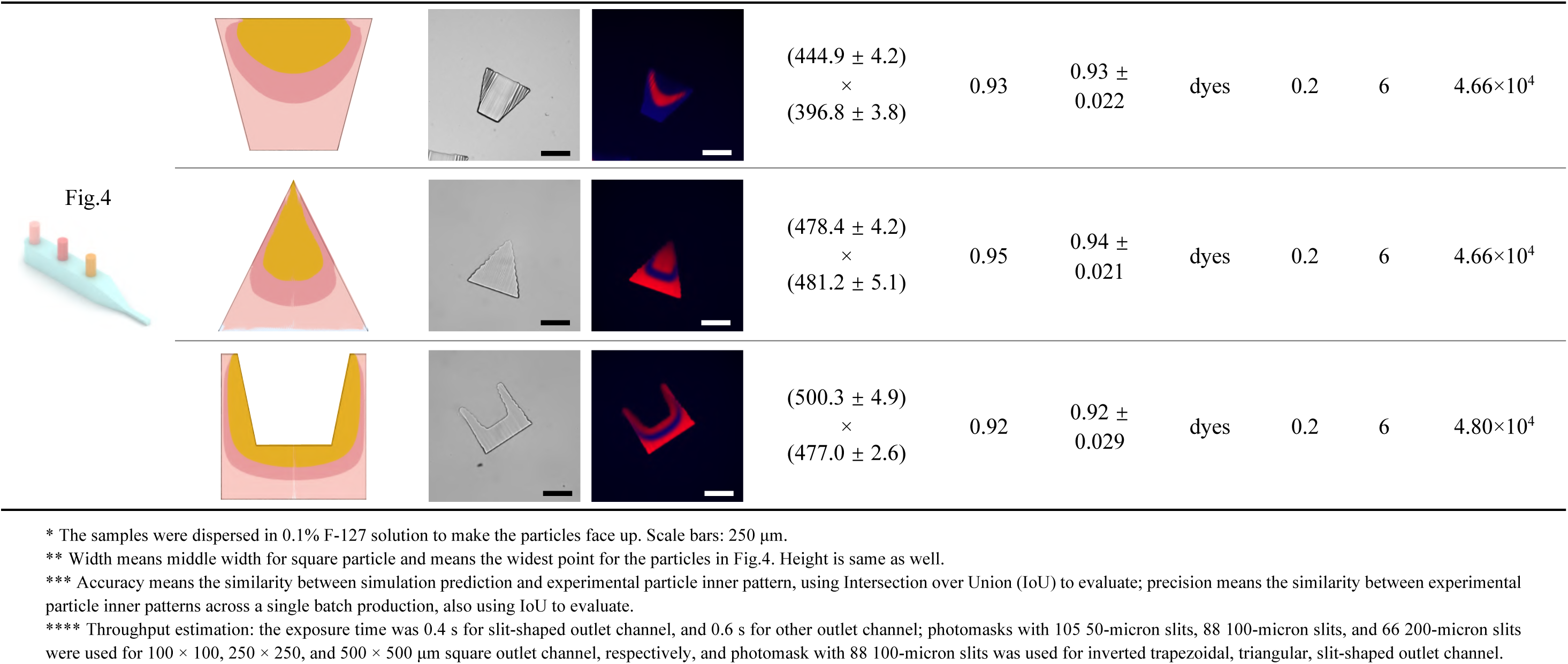
Parameters for patterned microparticles fabricated with multi-inlet configurations.

**Table S5.**
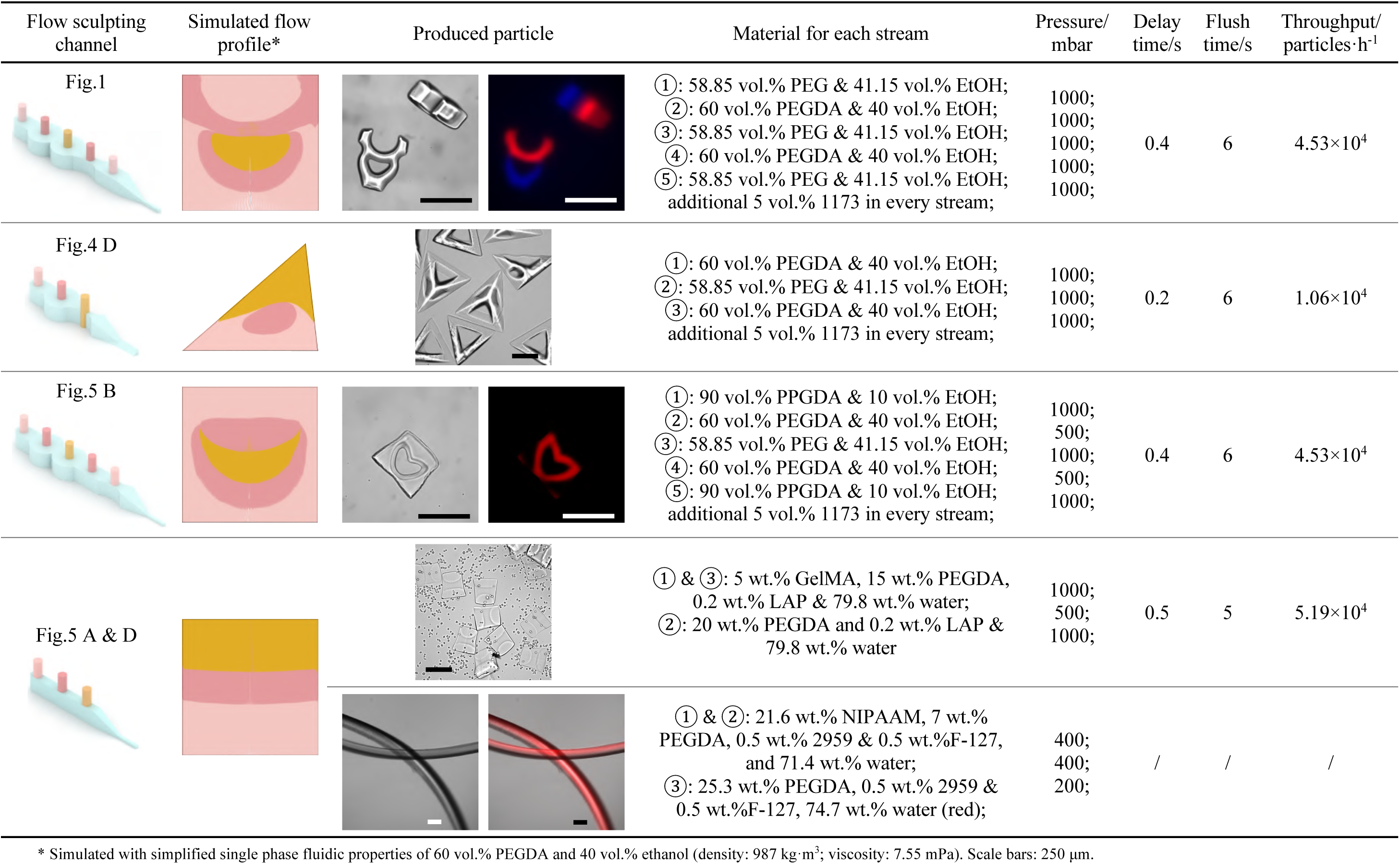
Parameters for structured multi-material microparticles fabricated with multi-inlet configurations.

| Flow sculpting channel | Simulated flow profile* | Produced particle | Material for each stream | Pressure/<br>mbar | Delay<br>time/s | Flush<br>time/s | Throughput/<br>particles·h<sup>-1</sup> |
| --- | --- | --- | --- | --- | --- | --- | --- |
| Fig.1 |  |  | ①: 58.85 vol.% PEG & 41.15 vol.% EtOH;<br>②: 60 vol.% PEGDA & 40 vol.% EtOH;<br>③: 58.85 vol.% PEG & 41.15 vol.% EtOH;<br>④: 60 vol.% PEGDA & 40 vol.% EtOH;<br>⑤: 58.85 vol.% PEG & 41.15 vol.% EtOH;<br>additional 5 vol.% 1173 in every stream; | 1000;<br>1000;<br>1000;<br>1000;<br>1000; | 0.4 | 6 | 4.53×10<sup>4</sup> |
| Fig.4 D |  |  | ①: 60 vol.% PEGDA & 40 vol.% EtOH;<br>②: 58.85 vol.% PEG & 41.15 vol.% EtOH;<br>③: 60 vol.% PEGDA & 40 vol.% EtOH;<br>additional 5 vol.% 1173 in every stream; | 1000;<br>1000;<br>1000; | 0.2 | 6 | 1.06×10<sup>4</sup> |
| Fig.5 B |  |  | ①: 90 vol.% PPGDA & 10 vol.% EtOH;<br>②: 60 vol.% PEGDA & 40 vol.% EtOH;<br>③: 58.85 vol.% PEG & 41.15 vol.% EtOH;<br>④: 60 vol.% PEGDA & 40 vol.% EtOH;<br>⑤: 90 vol.% PPGDA & 10 vol.% EtOH;<br>additional 5 vol.% 1173 in every stream; | 1000;<br>500;<br>1000;<br>500;<br>1000; | 0.4 | 6 | 4.53×10<sup>4</sup> |
| Fig.5 A & D |  |  | ① & ③: 5 wt.% GelMA, 15 wt.% PEGDA,<br>0.2 wt.% LAP & 79.8 wt.% water;<br>②: 20 wt.% PEGDA and 0.2 wt.% LAP &<br>79.8 wt.% water | 1000;<br>500;<br>1000; | 0.5 | 5 | 5.19×10<sup>4</sup> |
|  |  |  | ① & ②: 21.6 wt.% NIPAAm, 7 wt.%<br>PEGDA, 0.5 wt.% 2959 & 0.5 wt.% F-127,<br>and 71.4 wt.% water;<br>③: 25.3 wt.% PEGDA, 0.5 wt.% 2959 &<br>0.5 wt.% F-127, 74.7 wt.% water (red); | 400;<br>400;<br>200; | / | / | / |

\* Simulated with simplified single phase fluidic properties of 60 vol.% PEGDA and 40 vol.% ethanol (density: 987 kg·m<sup>-3</sup>; viscosity: 7.55 mPa). Scale bars: 250 μm.

**Figure S1.**
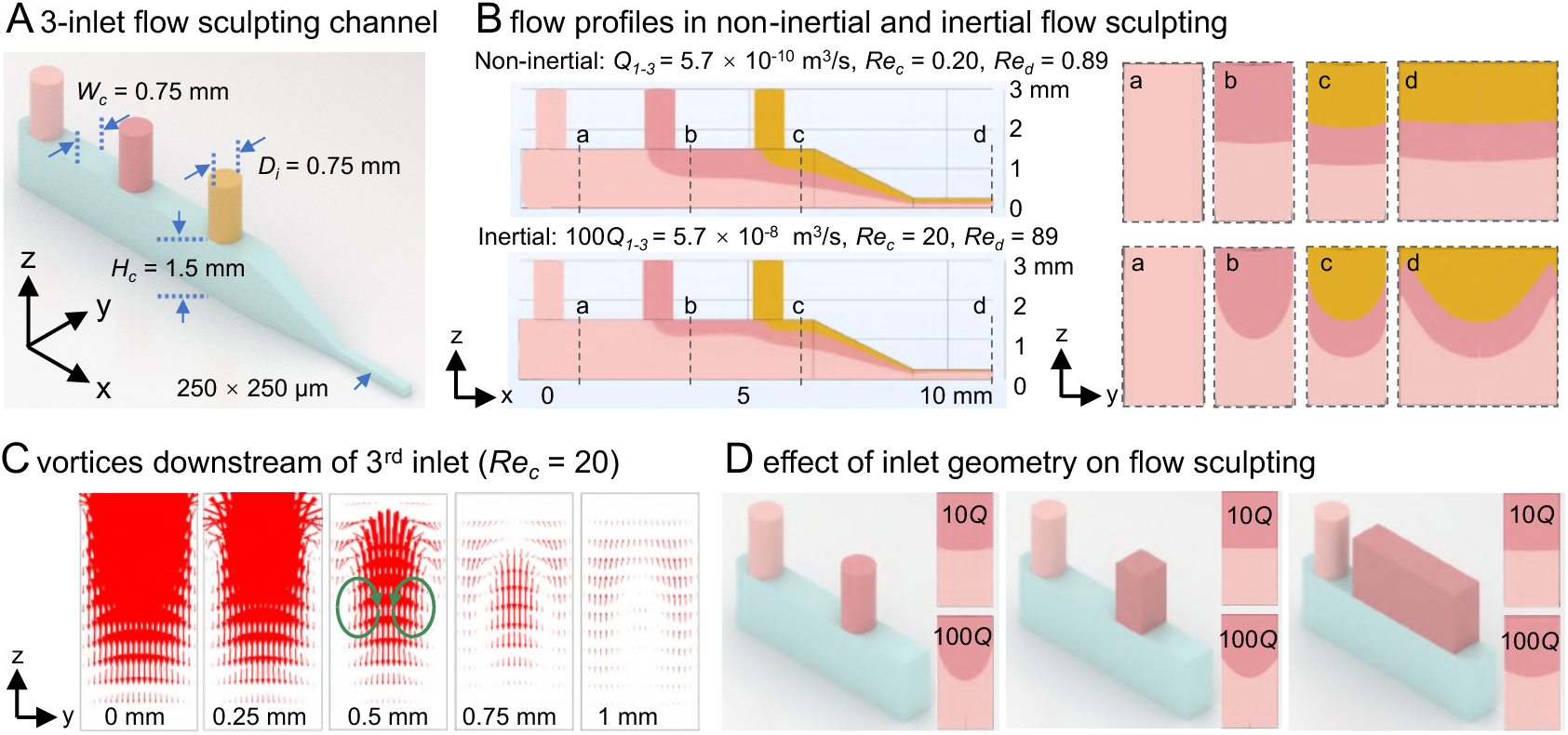
Non-inertial vs. inertial flow sculpting. (A) A representative flow sculpting microchannel with three inlets arranged in series following inlet configuration one (Σ*D_i_* = *W_c_*). (B) Numerical simulations model the multi-layered flow profile for non-inertial, *Re* ≲ *O*(1), and inertial, *Re* ≈ *O*(10), flow regimes. (C) Dean flow vortices produced downstream of the 3^rd^ inlet due to inertial flow. The vector plots are obtained from YZ cross-sectional planes at distances 0, 0.25, 0.5, 0.75, and 1mm downstream of the 3^rd^ inlet. (D) Circular, squared, and rectangular inlet geometries produce different flow profiles at varying *Re*. **Note S1. Non-inertial vs. inertial flow sculpting.** Three inlets (*D_i_* = 0.75 mm) are arranged in series along a rectangular cross-sectional (*W_c_* = 0.75 mm, *H_c_* = 1.5 mm) segment of a flow sculpting channel following inlet configuration one, i.e., Σ*D_i_* = *W_c_*, (Figure S1A). The rectangular channel converges into a square outlet channel (250 × 250 µm) to obtain the final sculpted flow profile. We perform numerical simulations to model the multi-layered flow profile for non-inertial, *Re* ≲ *O*(1), and inertial, *Re* ≈ *O*(10), flow regimes (Figure S1B). The non-inertial flow with *Re_c_* = 0.20 and *Re_d_* = 0.89 within the rectangular and squared channel cross-section, respectively, resulted in a vertical and flat stacking of the three layers at the outlet. However, the inertial flow with 100x higher flow rate, i.e., *Re_c_* = 20 and *Re_d_* = 89, produced a downward curvature in the three stacked layers, which is attributed to the Dean flow vortices produced by the high velocity flow from the inlet turning by 90 degrees into the main channel. A vector plot within the rectangular cross-section clearly indicates the presence of the Dean flow vortices produced downstream of the 3^rd^ inlet due to inertial flow, which gradually disappears away from the inlet (Figure S1C). This inertial flow effect and subsequent Dean vortices could be minimized by altering the shape and size of the inlet channel (Figure S1D). As the inlet geometries are modified from circular to square and rectangular cross-sections, with longer dimension along the flow direction, the curvature effect is reduced gradually even at higher *Re*. A wider rectangular inlet port will produce a relatively lower flow velocity for the given flow rate and will allow a longer time for the two merging streams to interact, thereby reducing the inertial effects and Dean flow vortices that curve the interface between the layers.

**Figure S2.**
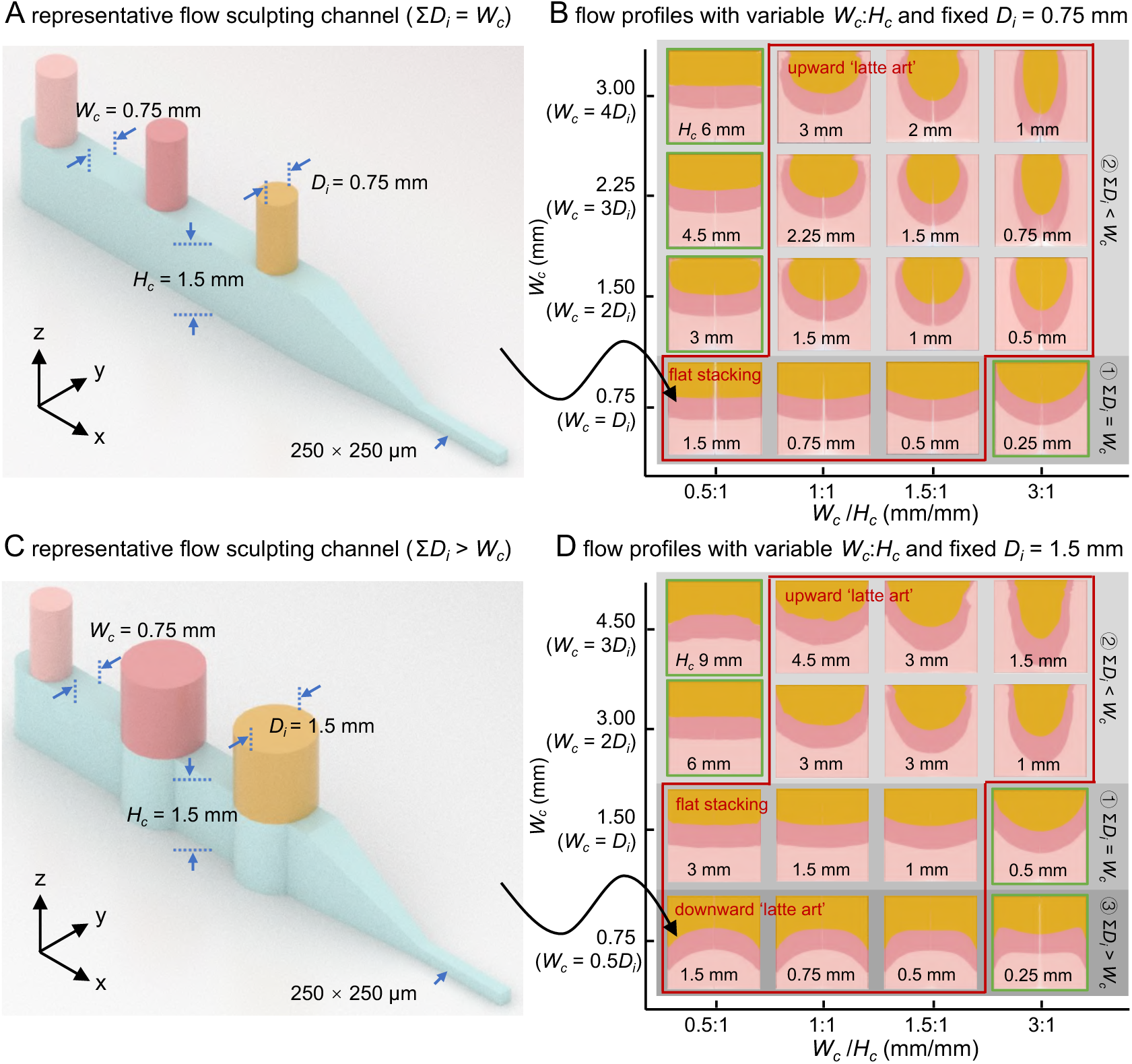
Sculpted flow profiles with variable inlet diameter (*D_i_*), channel width (*W_c_*), height (*H_c_*), and aspect ratio (*W_c_*/*H_c_*). (A) A representative flow sculpting microchannel with three inlets arranged in series using the inlet configuration one (Σ*D_i_* = *W_c_*). (B) Sculpted flow profiles at the outlet following different inlet configurations, i.e., Σ*D_i_* = *W_c_*, and Σ*D_i_* < *W_c_*, using a constant *D_i_* of 0.75 mm. *W_c_* is varied from 0.75 mm to 3 mm. *H_c_* is varied from 0.25 mm to 6 mm. (C) A representative flow sculpting microchannel with three inlets arranged in series using the inlet configuration three (Σ*D_i_* > *W_c_*). (D) Sculpted flow profiles at the outlet following different inlet configurations, i.e., Σ*D_i_* = *W_c_*, Σ*D_i_* < *W_c_*, and Σ*D_i_* > *W_c_*, using a constant *D_i_* of 1.5 mm. *W_c_* is varied from 0.75 mm to 4.5 mm. *H_c_* is varied from 0.25 mm to 9 mm. Green boxes: unexpected deviation, red boxes: expected flow profiles. **Note S2. Sculpted flow profiles with variable inlet diameter (*D_i_*), channel width (*W_c_*), height (*H_c_*), and aspect ratio (*W_c_*/*H_c_*).** The shape of the sculpted flow profiles for the three inlet configurations (Σ*D_i_* = *W_c_*, Σ*D_i_* < *W_c_*, and Σ*D_i_* > *W_c_*) holds consistently for a broad range of the microchannel dimensions (*D_i_*, *W_c_*, *H_c_*, and *W_c_*/*H_c_*) (Figure S2). However, deviation from the expected flow profile shapes was observed for limited cases (green boxes). For example, a vertical and flat stacking of three flow streams, following inlet configuration one (Σ*D_i_* = *W_c_,* the fourth row of Figure S2B and the third row of Figure S2D), was as expected under *W_c_*/*H_c_* ≤ 1.5, whereas, a higher aspect ratio channel with *W_c_*/*H_c_* = 3 resulted in enhanced upward curvature (green boxes in the fourth row of Figure S2B and the third row of Figure S2D) mimicking the inlet configuration two. For the inlet configuration two (Σ*D_i_* < *W_c_*, the first-third rows of Figure S2B and the first-second rows of Figure S2D), the stacked layers are curved upwards as expected for *W_c_*/*H_c_* ≥ 1 (red boxes); however, for *W_c_*/*H_c_* = 0.5, we observed unexpectedly vertical stacking of the layers mimicking the inlet configuration one (green box). For the inlet configuration three (Σ*D_i_* > *W_c_*, the last row of Figure S2D), we obtained a sculpted flow profile with stacked layers curved downwards as expected for *W_c_*/*H_c_* ≤ 1.5, whereas, for *W_c_*/*H_c_* = 3, the flow profile with reduced curvature deviates slightly from the expected shape.

**Figure S3.**
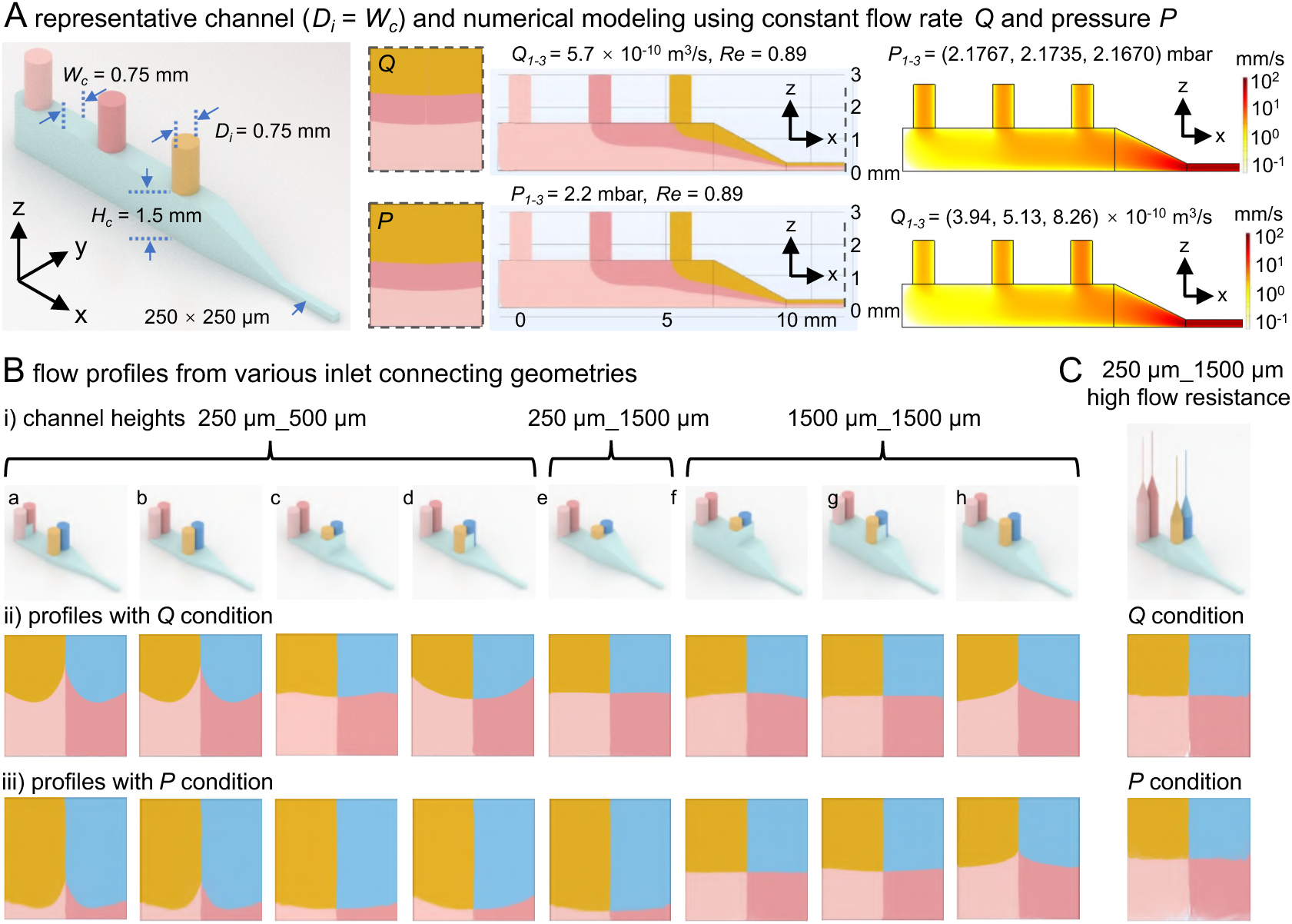
Defining appropriate inlet boundary conditions in the numerical models: flow rate *Q* vs. pressure *P*. (A) A representative flow sculpting microchannel with inlet configuration one (Σ*D_i_* = *W_c_*), and numerical modeling of sculpted flow profile using inlet boundary conditions based on constant flow rate *Q* and pressure *P* across three inlets within non-inertial flow regime, i.e., *Re* ≲ *O*(1). (B) Comparison of sculpted flow profiles resulting from different channel designs and inlet connections under constant *Q* and *P* boundary conditions. (C) A microchannel model mimicking high flow resistances across four inlets results in similar sculpted flow profiles for constant *Q* and *P* boundary conditions. **Note S3. Defining appropriate inlet boundary conditions in the numerical models: flow rate *Q* vs. pressure *P*.** We conducted numerical simulations using a 3-inlet flow sculpting microchannel (Σ*D_i_* = *W_c_*) to investigate the effects of inlet boundary conditions (BCs) with constant flow rate *Q* of 5.7×10^-10^ m^3^/s or pressure *P* of 2.2 mbar (corresponding to an outlet *Re* of 0.89) on the resulting tri-layered vertically stacked flow profiles and velocity distributions (Figure S3A). A constant *Q* boundary condition with flow rate ratio *Q_1_*:*Q_2_*:*Q_3_* = 1:1:1 resulted in layer thicknesses with a symmetric ratio of *t_1_*:*t_2_*:*t_3_* ≈ 1.5:1:1.5 (i.e., 37.5%, 25%, and 37.5% of the channel height) defined by the Poiseuille velocity profile. For a constant *Q* boundary condition, the first inlet converges numerically with the highest inlet pressure of 2.1767 mbar, followed by 2.1735 and 2.1670 mbar for the second and third inlets, respectively. The inlet pressures vary across the channel length; however, the relative proportions of different stream layers remained symmetric following the Poiseuille velocity profile and constant *Q*. In comparison, a constant *P* (= 2.2 mbar, similar to the converged pressure values from the last numerical model) boundary condition with pressure ratio *P_1_:P_2_:P_3_* = 1:1:1 resulted in layer thicknesses with an asymmetric ratio of *t_1_*:*t_2_*:*t_3_* ≈ 1.29:1:2.18 (i.e., 28.8 %, 22.4 %, and 48.8 % of the channel height). Different hydraulic flow resistances (*R_h_*) for a constant pressure drop *ΔP* = 2.2 mbar between the three inlets and the outlet translated into variable flow rates of 3.94, 5.13, and 8.26 ×10^-10^ m^3^/s, respectively (following Hagen-Poiseuille equation: *Q* = Δ*P*/*R_h_*). Hence, variable layer thicknesses within the sculpted flow profile were obtained, which were consistent with the inlet flow rates. We have further investigated various inlet designs with different flow resistances for inlets positioned along the channel length (Figure S3B). The connecting geometries between two juxtaposed inlets (parallel configuration) affect the interface between the co-flowing streams from these inlets and the final sculpted flow profile. Notably, shallower channels (*H_c_* = 250 µm between the series inlets), with a greater difference in *R_h_* between inlets placed in series along the channel length, produce flow profiles having thinner bottom layers associated with the first two inlets for a constant *P* boundary condition. For a constant *Q* boundary condition, the layer thicknesses within the checkered flow profile are relatively consistent for different channel designs; however, the interface shape is influenced by the inter-inlets geometric feature. Once the channel height is increased (*H_c_* = 1,500 µm between the series inlets), the difference in *R_h_* between inlets along the channel length is reduced. The bottom layer thickness for the constant *P* boundary condition starts to approach that for the constant *Q* boundary condition. We mitigate the inter-inlets difference in *R_h_* by introducing additional flow resistance into the system (mimicking connecting tubing with much smaller diameters), such that the sculpted flow profile is not affected by the minor flow resistance variation between the inlets (Figure S3C). The flow profile obtained for constant *Q* and *P* boundary conditions matched well. Therefore, we have adopted these findings in our experimental workflow by using inlet connecting tubing with significantly higher flow resistances compared to the channel itself.

**Figure S4.**
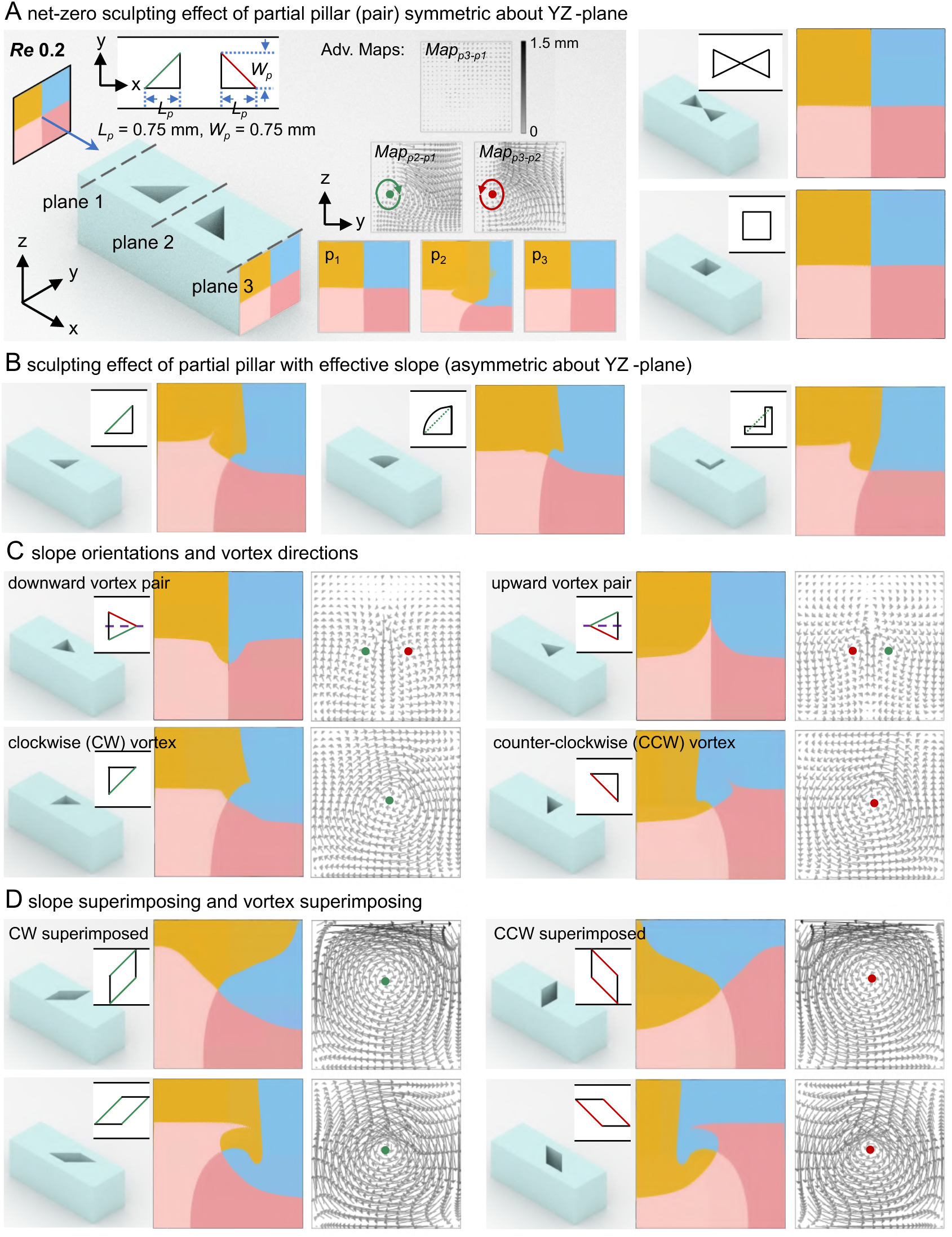
Flow sculpting principle of a partial-height pillar in non-inertial flow. (A) A representative flow sculpting mode showing the input (initial flow profile from a given inlet configuration), unit operator (partial-height pillar structure), and output (resulting flow profile). Any partial-height pillar or pillar pair structure symmetric about the YZ-plane produces a net-zero sculpting effect on the flow profile. Advection maps indicate the direct flow transformation (advection vortices) between specific planes. (B) Effective slopes in the partial-height pillars are responsible for inducing the sculpting effect. The direction (C) and the intensity (D) of these advection vortices are determined by the orientation and the superimposition of these effective slopes. **Note S4. Flow sculpting principle of a partial-height pillar in non-inertial flow.** Flow profile reshaping in the YZ plane is achieved by generating secondary flows in this plane, which are perpendicular to the direction of primary flow. Such flow reshaping can occur either through Dean flow in the inertial flow regime or through flow dislocation in the non-inertial flow regime. Since the flow must remain continuous, any local fluid displacement would be immediately compensated by the surrounding fluid. Consequently, changes in the flow profile usually manifest as 2D vortex-like flow transformations within the plane. In non-inertial flow sculpting, this type of flow transformation, i.e., flow dislocation, is initially induced by the lateral offsets (y-direction) between flow layers obstructed by pillar structures and those that remain unobstructed as the fluid moves downstream the primary flow direction. The transformation is then completed by forming advection vortices under fluid advection. The oblique sides of partial-height pillar structures serve to generate these lateral offsets. Any given initial flow profile will be mapped to a corresponding output profile after passing through a partial-height pillar with an oblique side. Advection maps reveal the direct flow transformations produced by specific partial-pillars. For example, *Map_p2-p1_* illustrates exactly how the fluid flows and is transformed through the first pillar, from the flow profile at plane 1 to the flow profile at plane 2. Comparing *Map_p2-p1_* and *Map_p3-p2_* shows that this YZ-plane-symmetric partial-height pillar pair generates pattern-matched, counter-rotating advection vortex pairs, resulting in a net-zero sculpting effect (Figure S4A,). In contrast, a YZ-plane-asymmetric partial-height pillar with effective slope can induce a non-net-zero flow sculpting effect (Figure S4B), where the direction and the intensity of these advection vortices are governed by the orientation (Figure S4C) and superposition (Figure S4D) of these effective slopes. Closer inspection reveals that the plane 2 flow profile of Figure S4A is not identical to the leftmost flow profile of Figure S4B, even though they share the same initial flow profile and pass through the same pillar structure. This difference arises from hydrodynamic interactions between adjacent pillars, which persist until the pillars are sufficiently spaced apart (Figure 3C).

**Figure S5.**
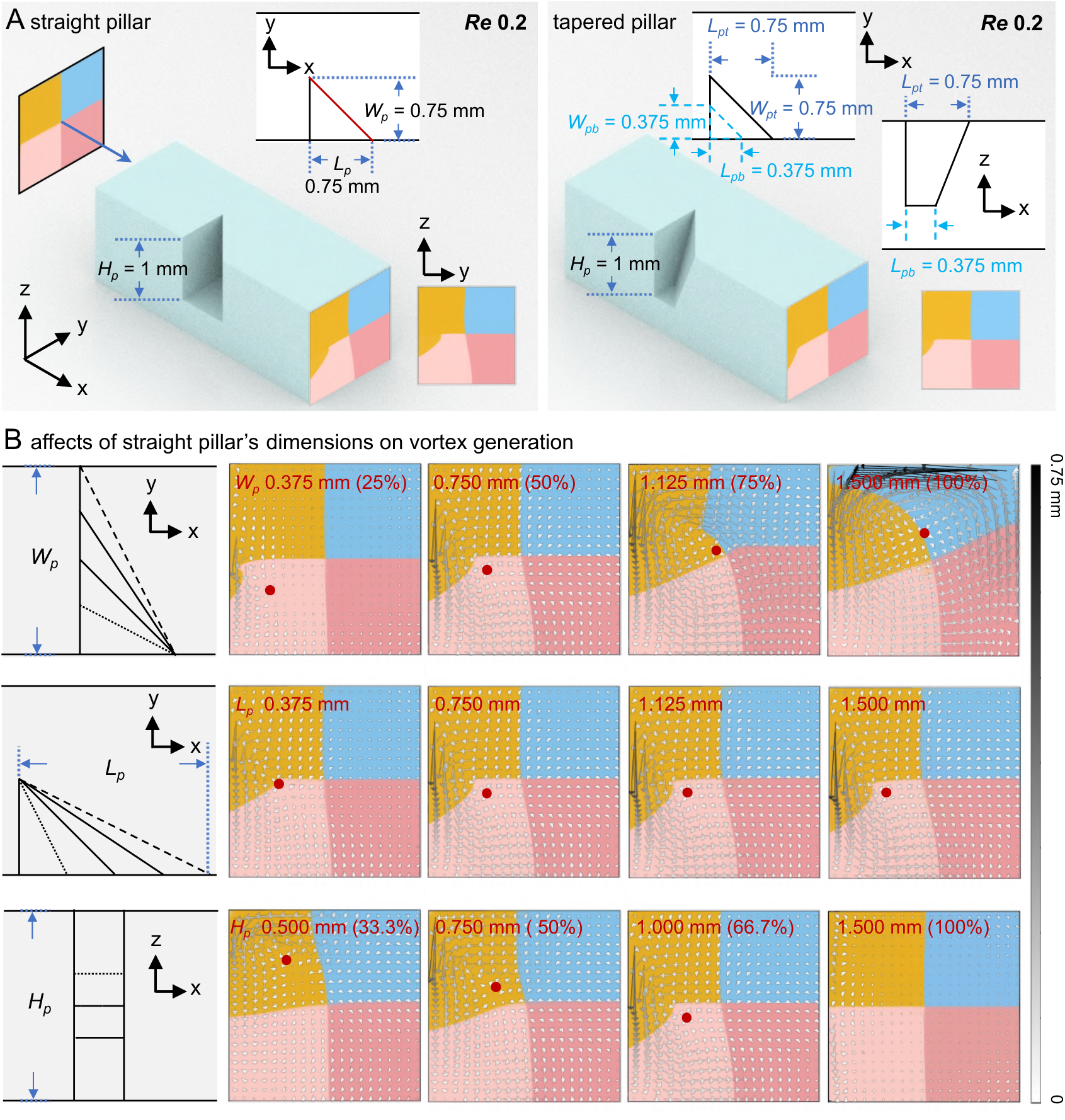
Effect of pillar dimensions on non-inertial flow sculpting. (A) Comparison of flow sculpting effects produced by straight vs. tapered pillars. The tapered pillar generates smaller lateral offsets in the y-direction compared to the straight pillar, thereby resulting in a gentler flow sculpting effect. (B) Influence of the straight pillar’s width (*W_p_*), length (*L_p_*), and height (*H_p_*) on the flow dislocation (advection maps showing the advection vortices) and, consequently, on the final flow sculpting effect (flow profiles). **Note S5. Effect of pillar dimensions on non-inertial flow sculpting.** In flow reshaping, the flow dislocation (advection vortex) observed on the YZ-plane represents the projection of the 3D vortex within the channel onto this plane. Consequently, the geometric dimensions of the pillar that project onto the YZ-plane, i.e., width (*W_p_*) and height (*H_p_*), significantly affect the central position and region of the resulting advection vortex on this plane, while the pillar length (*Lₚ*) exerts only a minor influence on this.

**Figure S6.**
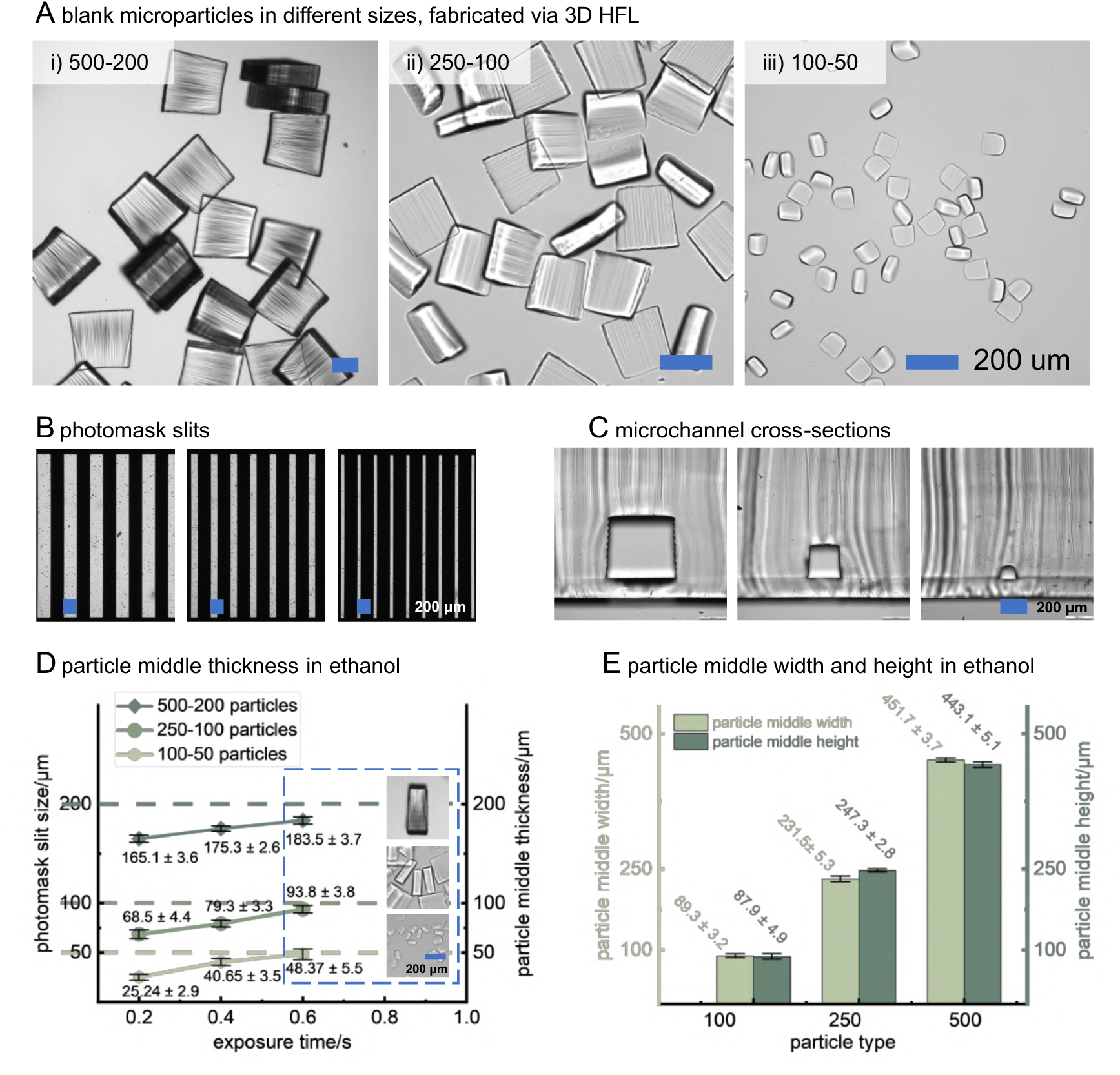
Thickness and cross-sectional size controls of square microparticles fabricated by 3D Hydrodynamic Flow Lithography method. (A) Microscope images of microparticle in large (∼500 μm), medium (∼250 μm) and small (∼100 μm) sizes. Particles were dispersed in ethanol after washing, to make as many sides face upwards. (B) Photomasks and (C) microchannels used for square microparticle production. (D) Middle thickness measurements of microparticle with different sizes and exposure times. (E) middle width and middle height measurements of microparticle with different sizes, 0.4 s and 0.6 s exposure time were used for 100 and 250/500 μm particle fabrication, respectively. Scale bars: 250 μm. **Note S6. Thickness and cross-sectional size controls of square microparticles fabricated by 3D Hydrodynamic Flow Lithography method.** When the exposure intensity is sufficient, the cross-sectional size of the particle is mainly determined by the cross-sectional size of the outlet channel, while the thickness of the particle is mainly determined by the slit pattern of the photomask. By varying the sizes of the channel and the photomask slit, particles of different sizes can be produced. The production throughput, measured in particles per hour, can be calculated based on the number of particles exposed per cycle by the photomask and the total cycle time, which includes exposure time *t*_exposure_, delay time *t*_delay_, and flush time *t*_flush_, as expressed by the formula:

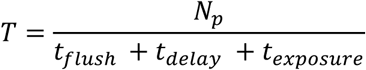

to roughly estimate the massive production capacity of the SFL setup in this work, we tested three types particles: one produced with the 100 μm channel and the 50 μm photomask slit (abbreviated as 100-50 to represent this type of particle), another with the 250 μm channel and the 100 μm photomask slit (250-100), and the third one with the 500 μm channel and the 200 μm photomask slit (500-200), representing small, medium, and large size particles, respectively. Their minimum exposure times, delay times, and flush times were studied and summarized with corresponding throughputs detailed in Tables S3-5. In general, longer exposure time allows the precursor solution to gain more polymerization within the restrictions from photomask slits and channel sizes (Figure S6B). However, due to the gradual absorption and attenuation of UV light with depth in the solution, the particles produced were usually not uniform in thickness and cross section, with the side closer to the light source being thicker and the side away from the light source being thinner. This disparity was more obvious in larger-sized particles. For instance, the middle thickness of 500-200 particles was only ∼ 180 μm after a 0.6 s exposure, even though the bottom reached 200 μm slit size. Increasing exposure time can alleviate this unevenness to some extent, but prolonged exposure time may lead to overexposure and connect the particles into fibers when the gap between slits is not sufficient (e.g. 0.8 s exposure time with 200-200 (slit-gap) photomask), namely, sufficient gap to slit ratio should be guaranteed for the adequate but not excessive polymerization under strong UV exposure.

**Figure S7.**
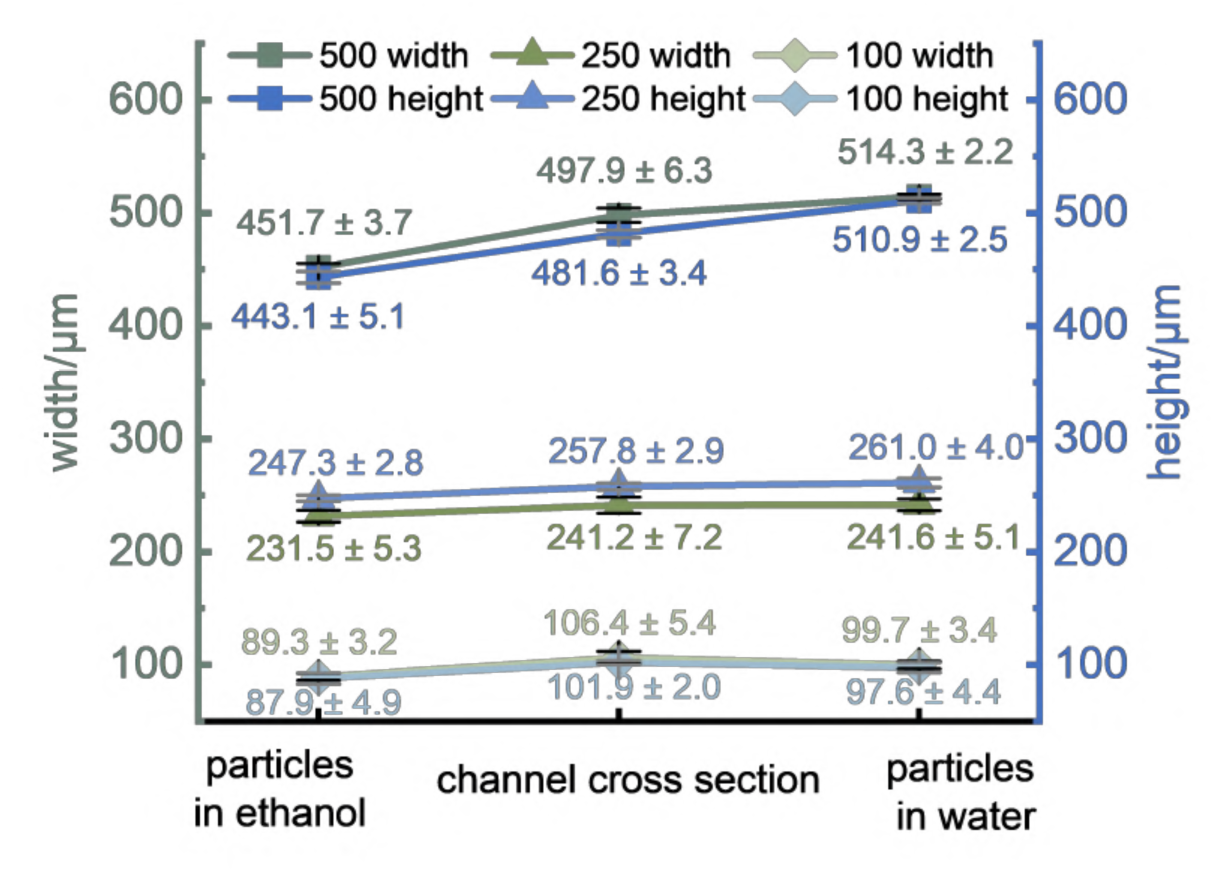
**Width and height measurements of microchannel cross section and fabricated square microparticles dispersed in ethanol and water.**

**Figure S8.**
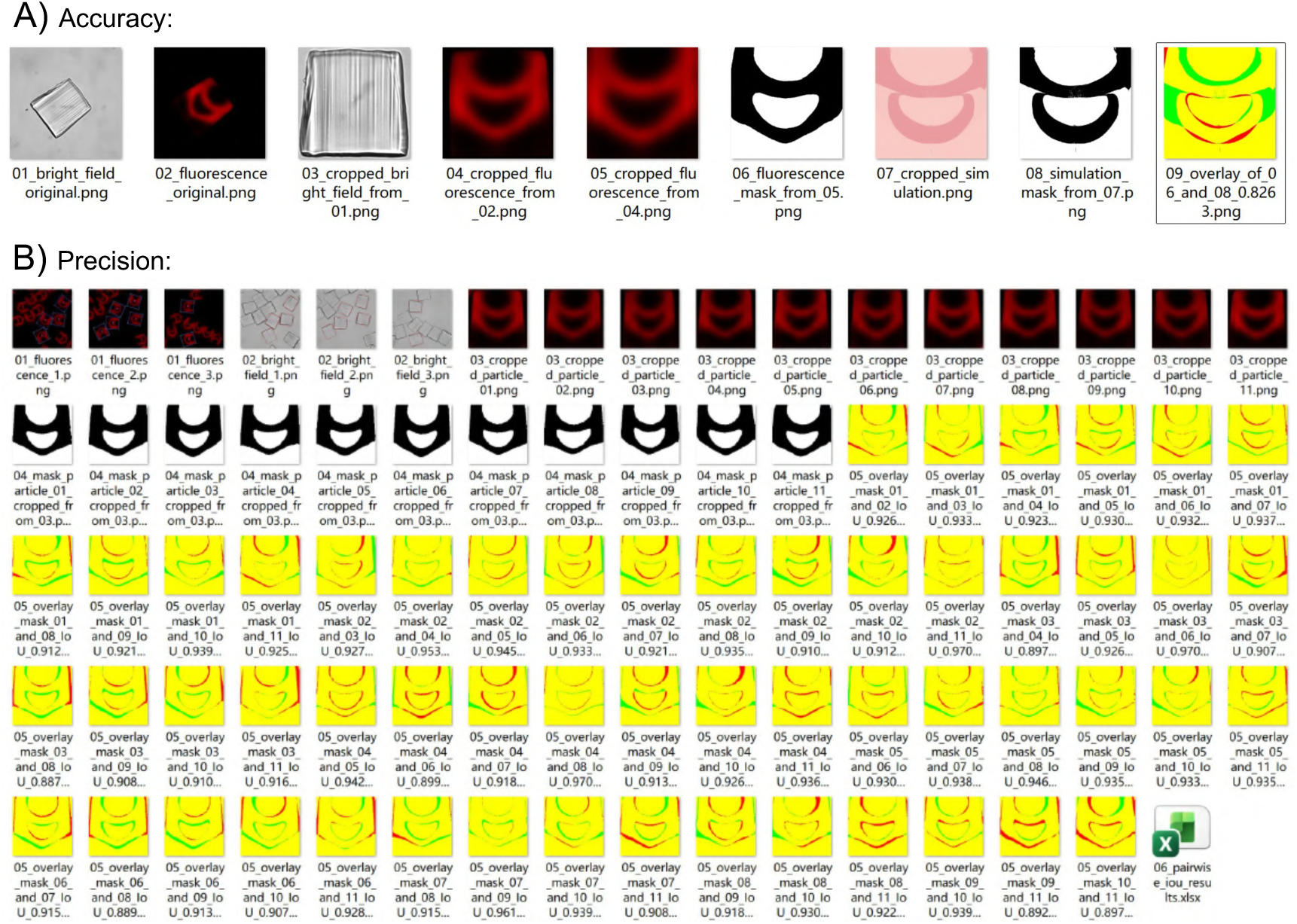
IoU estimations for the particle pattern accuracy and precision. (A) Accuracy: similarity between simulation prediction and experimental particle inner pattern. In image_09, red shows simulation-only regions, green shows experiment-only regions, and yellow indicates regions where both overlap (IoU, 0.83). (B) Precision: similarity between experimental particle inner patterns. In overlay images, red shows particle_a-only regions, green shows particle_b-only regions, and yellow indicates regions where both overlaps. Mean IoU, 0.92 ± 0.022, was estimated by the pairwise IoUs from multiple random particles across a single batch production. All image pre-processing and IoU computation were carried out using custom Python scripts.

**Figure S9.**
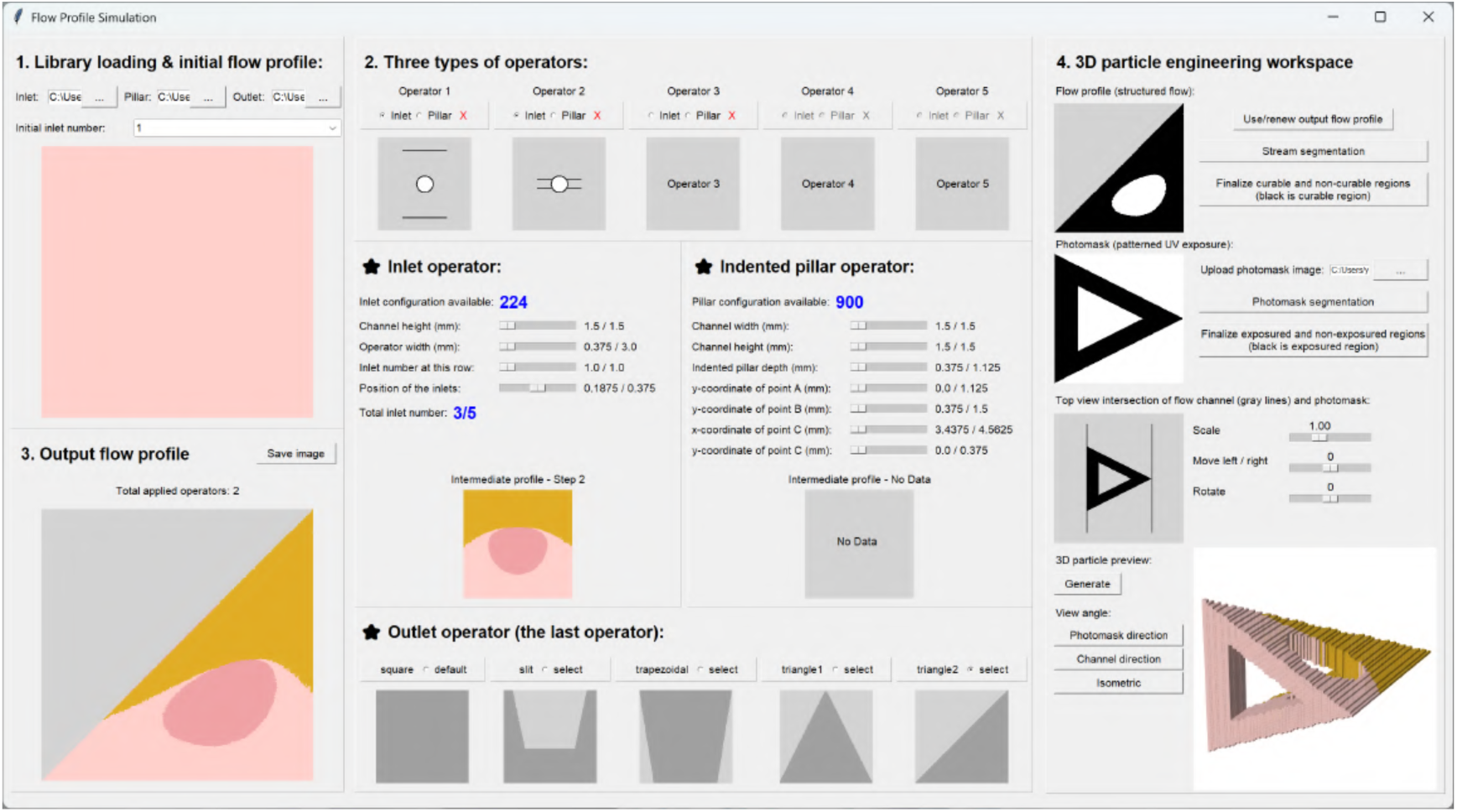
Representative GUI design parameters used to generate the sculpted flow profile and 3D microparticles presented in. **Figure 4D**.

**Figure S10.**
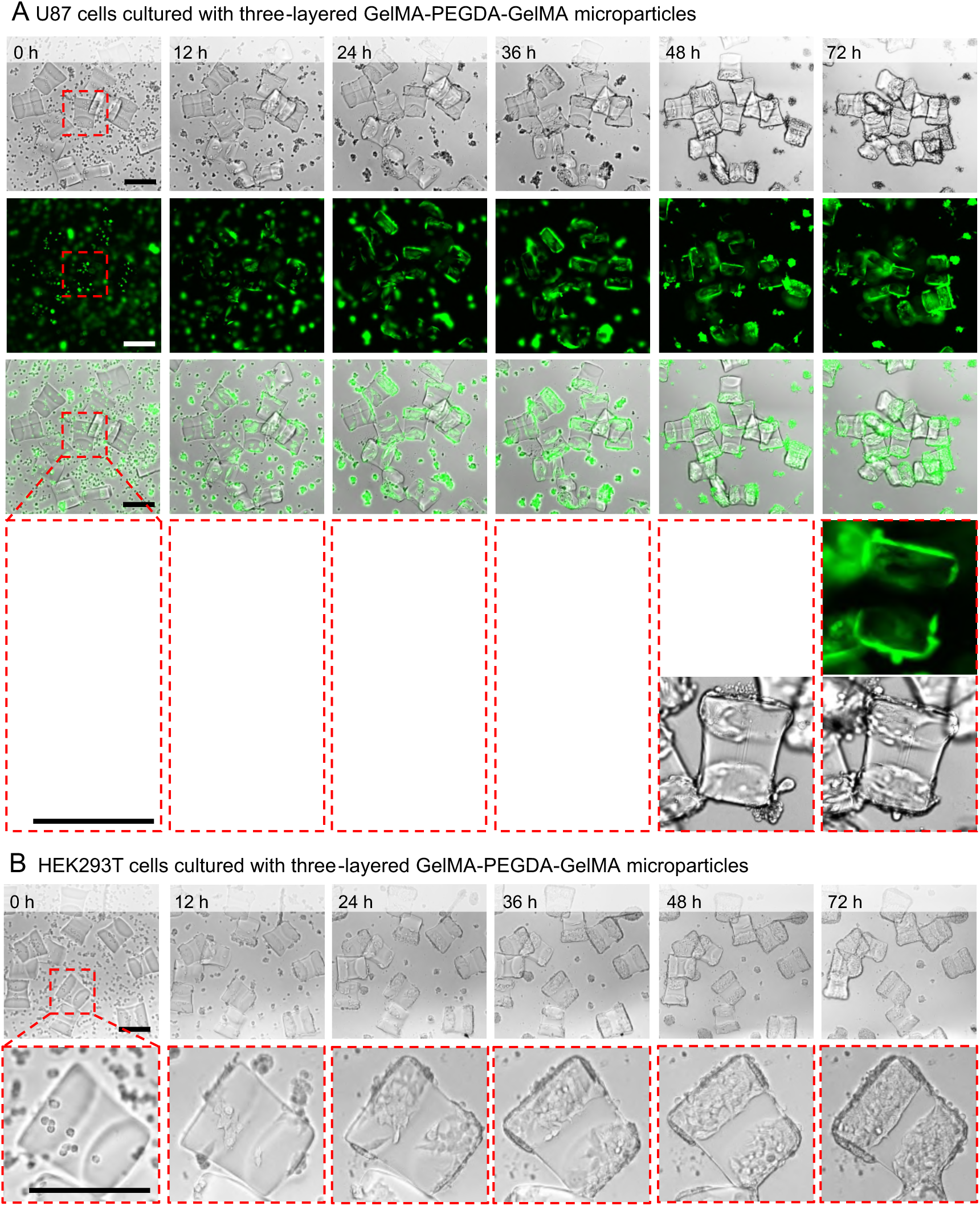
Time-lapse images of different cell types cultured with three-layered GelMA-PEGDA-GelMA microparticles. (A) GFP-expressing U87 glioblastoma cell line. From top to bottom: bright field, fluorescent, overlapped, and zoomed-in fluorescent and bright field images. From left to right: images captured after 0 h, 12 h, 24 h, 36 h, 48 h, and 72 h of culture. (B) human embryonic kidney cell line HEK293T. Top: bright-field images; bottom: zoomed-in bright-field images. From left to right: images captured after 0 h, 12 h, 24 h, 36 h, 48 h, and 72 h of culture. Cells migrate gradually towards the particles and adhere exclusively to the top and vertical surfaces of the GelMA region, avoiding the PEGDA area. Scale bars: 250 μm.

Movie 1 Fast flow profile engineering and 3D particle visualization via GUI.

Movie 2 Culture of different cell types with three-layered GelMA–PEGDA–GelMA particles.

